# *Fusobacterium nucleatum* determines the expression of amphetamine-induced behavioral responses through an epigenetic phenomenon

**DOI:** 10.1101/2025.01.15.633210

**Authors:** Samuel J. Mabry, Xixi Cao, Yanqi Zhu, Caleb Rowe, Shalin Patel, Camila González-Arancibia, Tiziana Romanazzi, David P. Saleeby, Anna Elam, Hui-Ting Lee, Serhat Turkmen, Shelby N. Lauzon, Cesar E. Hernandez, HaoSheng Sun, Hui Wu, Angela M. Carter, Aurelio Galli

## Abstract

Amphetamines (AMPHs) are psychostimulants commonly used for the treatment of neuropsychiatric disorders. They are also misused (AMPH use disorder; AUD), with devastating outcomes. Recent studies have implicated dysbiosis in the pathogenesis of AUD. However, the mechanistic roles of microbes in AUD are unknown. *Fusobacterium nucleatum* (*Fn*) is a bacterium that increases in abundance in both rats and humans upon AMPH exposure. *Fn* releases short-chain fatty acids (SCFAs), bacterial byproducts thought to play a fundamental role in the gut-brain axis as well as the pathogenesis of AUD. We demonstrate that in gnotobiotic *Drosophila melanogaster,* colonization with *Fn* or dietary supplementation of the SCFA butyrate, a potent inhibitor of histone deacetylases (HDACs), enhances the psychomotor and rewarding properties of AMPH as well as its ability to promote male sexual motivation. Furthermore, solely HDAC1 RNAi targeted inhibition recapitulates these enhancements, pointing to a specific process underlying this *Fn* phenomenon. Of note is that the expression of these AMPH behaviors is determined by the increase in extracellular dopamine (DA) levels that result from AMPH-induced reversal of DA transporter (DAT) function, termed non-vesicular DA release (NVDR). The magnitude of AMPH-induced NVDR is dictated, at least in part, by DAT expression levels. Consistent with our behavioral data, we show that *Fn*, butyrate, and HDAC1 inhibition enhance NVDR by elevating DAT expression. Thus, the participation of *Fn* in AUD stems from its ability to release butyrate and inhibit HDAC1. These data offer a microbial target and probiotic interventions for AUD treatment.

## Introduction

Amphetamines (AMPHs) are psychostimulants commonly prescribed for the treatment of neuropsychiatric disorders such as attention deficit hyperactivity disorder^1^. AMPHs are also commonly misused, leading to devastating medical and societal effects. In the U.S. drug overdose deaths involving psychostimulants, primarily AMPHs, increased from 5,716 deaths in 2015 to an astonishing 34,022 deaths in 2021 (CDC WONDER). Yet, no pharmacological treatments are currently available. Multiple studies demonstrate that a robust communication between the gut microbiota and the brain is essential for healthy brain function, dopaminergic signaling, as well as animal behaviors^2,3^. Accordingly, imbalances in the gut microbiome (dysbiosis) have been implicated in the etiology and development of pathological behaviors such as psychostimulant use disorders (PUD)^4–7^. In support of this concept, antibiotic treatment reduces the psychomotor and rewarding effects of AMPHs, as well as AMPH-induced sensitization and reinstatement^7–13^. Interestingly, dysbiosis is observed in humans affected by AMPH use disorder (AUD)^14–17^. Of note is that an elevation in abundance of the opportunistic pathogen *Fusobacterium nucleatum* (*Fn*) is observed in AUD patients as well as rodents exposed to AMPH^15–17^. Heightened levels of this bacterium and others are thought to underlie multiple periodontal issues that are associated with AUD^18–20^. However, whether changes in the microbiome dictate the expression of behavioral responses to AMPH has yet to be uncovered.

Short-chain fatty acids (SCFAs), byproducts of microbial carbohydrate fermentation, are thought to play a critical role in the gut-brain axis and possess the ability to regulate neuronal processes^21,22^. Butyrate, acetate, and propionate are the main SCFAs produced by gut microbes^23^. Butyrate is known to cross the blood brain barrier^24,25^ and to enhance AMPH associated behaviors^26–28^. Yet, whether and how changes in the microbiota (*e.g. Fn* abundance) and its byproducts, including butyrate, participate in AUD remain elusive.

The psychomotor properties and the abuse potential of AMPHs have been associated with an increase in dopamine (DA) neurotransmission^29,30^. This increase stems, at least in part, from the ability of AMPH to enhance extracellular DA levels by promoting reverse transport of DA (<u>n</u>on-<u>v</u>esicular <u>D</u>A release; NVDR) *via* the DA transporter (DAT). The magnitude of AMPH-induced NVDR is dictated, at least in part, by DAT expression levels^31^. Furthermore, inhibition of NVDR reduces the ability of AMPH to exert its psychostimulant behavioral properties including preference and sexual motivation^32–36^. Thus, it is possible that changes in *Fn* abundance and its byproducts (*i.e.* butyrate) support AMPH psychostimulant properties by enhancing DAT expression and, consequently, NVDR.

As part of its molecular actions, AMPH causes profound changes in histone (H) acetylation^28,37^, an increasingly recognized epigenetic process that regulates gene transcriptional activation through chromatin remodeling^38,39^. Accordingly, AMPH leads to changes in gene transcriptional profiles^40–43^ required for long-term brain neuroadaptations^43^. The acetylation state of histones is determined by the action of two families of enzymes: histone acetyltransferase (HAT), which catalyzes the transfer of an acetyl group onto the histone tail, and histone deacetylase (HDAC), which catalyzes the removal of these acetyl groups. Consistent with the concept that *Fn* abundance supports the expression of AUD, butyrate, a HDAC inhibitor released by *Fn*^44,45^, leads both to an increase in histone acetylation^26^ and to changes in gene transcription profiles^46,47^ ^48,49^. Importantly, butyrate has also been shown to augment DAT expression^48,49^, a process associated with enrichment of acetylation of K9 and K14 of histone 3 (H3) within the DAT promotor^49^, and to enhance AMPH-induced locomotor sensitization^26–28^. This enhancement is consistent with the observation that an increase in DAT expression augments AMPH-induced NVDR as well as AMPH psychomotor properties^31,50^.

These data strongly suggest a potential role for *Fn* and butyrate in AUD and point to the targeting of specific bacterial populations as a possible pharmacotherapy. Consistent with this idea, the antibiotic minocycline, which decreases *Fn* counts in oral cavities of periodontitis patients^51^ also reduces the rewarding properties of AMPH both in humans and rodents^8,9^. Furthermore, the antibiotic ceftriaxone, that among others bacteria targets *Fn*^52^ also alters AMPH behavioral responses^12,13^.

In this study, we mechanistically elucidate how *Fn* enhances the expression of specific AMPH-associated behaviors including sexual motivation and drug preference. We demonstrate that *Fn*, through butyrate secretion, promotes DAT expression *via* selective inhibition of HDAC1. Increased DAT expression leads to enhanced NVDR, a process we show drives AMPH psychostimulant properties. These data strongly suggest a potential role for *Fn* and butyrate in AUD and point to the targeting of specific bacterial populations as a possible pharmacotherapy.

## Results

### *Fn* enhances both AMPH-induced psychomotor properties and NVDR

While recent evidence establishes a strong relationship between changes in the gut microbiome, altered dopaminergic signaling, and responses to drugs of abuse^2,4^, the specific microbes and mechanisms supporting this relationship have remained elusive^53^. Interestingly, *Fn* abundance is increased in humans affected by AUD^15–17^ and all antibiotics reported in the literature to decrease responses to AMPH target multiple species of fusobacteria, among others^8,9^. Therefore, we wanted to determine if *Fn* alone could alter the psychomotor and rewarding actions of AMPH. First, we rendered *Drosophilae* gnotobiotic by antibiotic (abx) treatment (see methods). Then, we colonized these flies by oral administration of *Fn* for 72 hours (Fig. 1A) in sucrose. We confirmed that bacteria derived from colonized flies matched characteristics of *Fn* by gram-stain (demonstrated gram-negative rods; Fig. 1A, right-top) and streaking on blood agar plates (demonstrated hemolytic activity; Fig. 1A, right-bottom). We found that *Fn* colonization increased AMPH-induced hyperlocomotion (Fig. 1B) a behavior strictly regulated by NVDR^32,33,36^. Consistently, *Fn* colonization also enhanced the ability of AMPH to promote NVDR, as measured by amperometry (Fig. 1C). The amperometric electrode was inserted between the PPL1 and PAM regions, two DA regions involved in startle-induced locomotion^54^ and drug preference^55^. These data demonstrate that *Fn* regulates AMPH actions.

**Figure 1:**
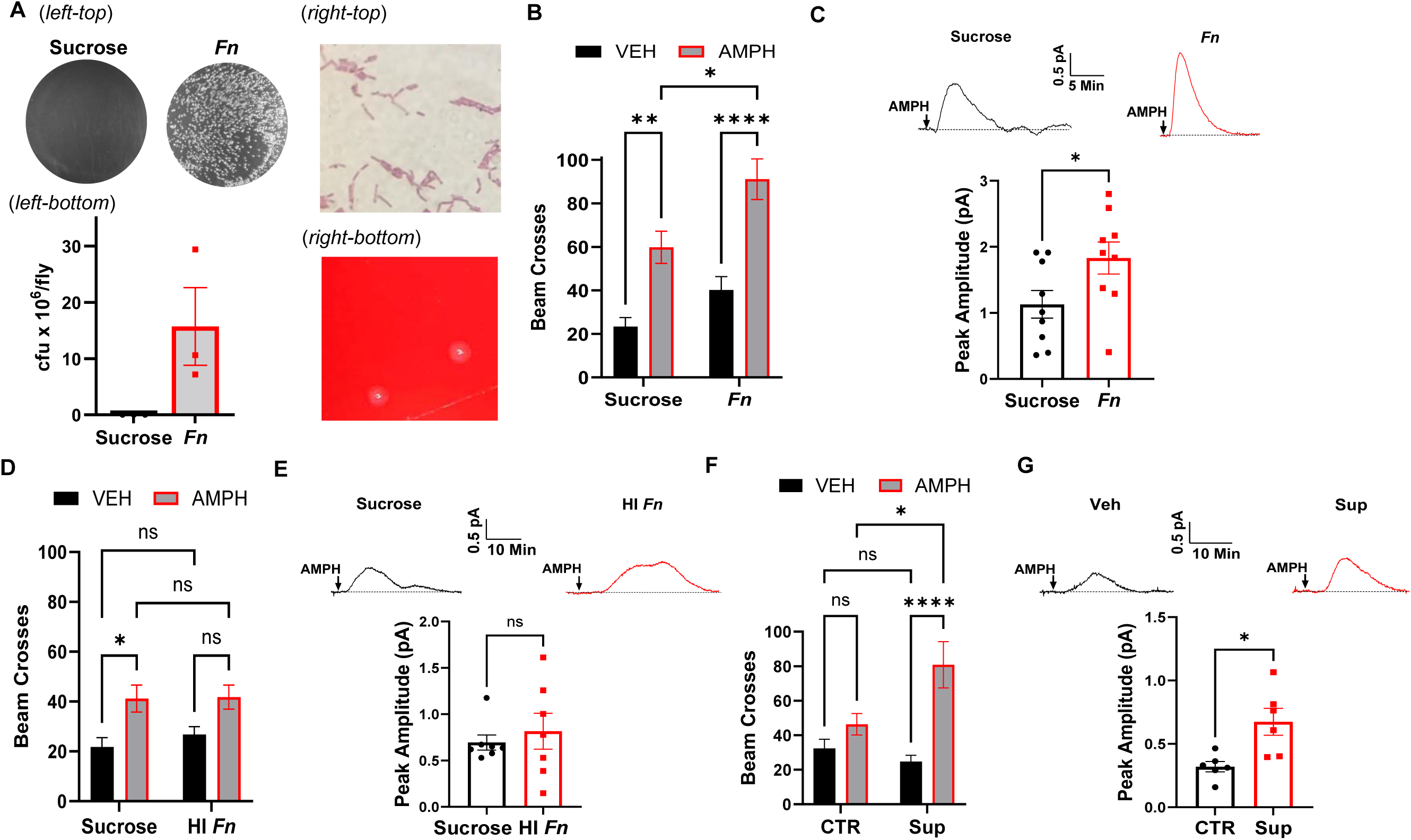
*Fn* enhances AMPH-induced hyperlocomotion and NVDR. All groups of *Drosophila* (Canton-S) were pretreated with abx (see methods) to reach gnotobiotic status. Flies were then orally administered specific treatments through capillary feeding for 72 h. Fly locomotor activity (panel B, D, F) was measured by beam crosses (100 min) in flies fed either vehicle (VEH, black bars) or AMPH (10 mM, grey bars). AMPH (20 µM)- induced NVDR (panel C, E, G) was determined by amperometry in isolated *Drosophila* brains. Representative traces are presented on top and quantitative analysis of the peak currents at the bottom of the panels. **A.** *Fn* fed flies were assessed for *Fn* colonization by homogenization and plating; representative images of plating (A, *left- top*) and overall cfu/fly (A, *left-bottom*). Characteristics of bacteria derived from flies were assessed by gram- stain (A, *right-top*) and plating on blood agar (A, *right-bottom*). **B.** AMPH induces hyperlocomotion in both sucrose and *Fn* treated flies. However, flies treated with *Fn* display a significant higher hyperlocomotion (F_1, 148_ = 11.91 for effect of *Fn*, p = 0.0007; F_1, 148_= 39.25 for effect of AMPH, p < 0.0001, n = 38-39). **C.** *Fn* colonization significantly enhances AMPH-induced NVDR (*t* = 2.191, p = 0.0436, n = 9). **D.** HI *Fn* does not enhance the ability of AMPH to cause hyperlocomotion (F_1, 178_ = 15.56 for effect of AMPH, p = 0.0001, n = 45-47). **E.** AMPH-induced NVDR is not significantly different in flies treated either with sucrose or HI *Fn* (U = 22, *p* = 0.8048, n = 7). **F.** AMPH-induced hyperlocomotion is increased in flies treated with the supernatant of the media used to grow *Fn* compared to flies treated with just media (F_1, 153_ = 6.888 for effect of interaction, p = 0.0096; F(1, 153) = 18.97 for effect of AMPH, p < 0.0001, n = 38-40). **G.** *Fn* supernatant treatment significantly enhances AMPH-induced NVDR (*t* = 3.124, p = 0.0108, n = 6). Data is presented as mean ± SEM. Two-way ANOVA with Tukey’s multiple comparison test (panels B, D, F); Student’s unpaired t-test (panels C, G); Mann-Whitney Test (panel E).

To determine whether stimulation of an immune response was involved, we fed flies heat inactivated (HI) *Fn*. *Fn* was autoclaved at 121 °C for 45 min. This metabolically inactive bacteria did not elicit any significant changes to either AMPH-induced hyperlocomotion (Fig. 1D) or NVDR (Fig. 1E). These observations suggested that *Fn* byproducts may be responsible for driving enhanced responses to AMPH. To determine if molecules secreted by *Fn* can enhance AMPH psychomotor properties, we orally treated flies with either unconditioned media or the supernatant obtained from growing cultures of *Fn*. Flies treated with the supernatant displayed enhanced psychomotor responses to AMPH (Fig. 1F) as well as elevated NVDR (Fig. 1G). To determine whether this ability to regulate AMPH actions is either specific to *Fn* or a general feature of gram-negative bacteria, we administered (as above) gnotobiotic flies either sucrose or sucrose containing VNP20009, an attenuated strain of *Salmonella enterica serovar typhimurium*. VNP20009 and have been engineered to be safe for *in vivo* administration by the stable knockout of msbB and purI genes^56,57^ and effectively colonize *Drosophila* as determined by plating of fly homogenates (Fig. S1A, left) and staining of fly gut tissue using antibodies specific for *Salmonella* (Fig. S1A, right). Importantly, colonization by VNP20009 did not significantly alter neither the ability of AMPH to cause hyperlocomotion (Fig. S1B) or NVDR (Fig. S1C), suggesting that this ability is not a general bacterial feature and may be specific to *Fn*.

### Butyrate and HDAC Inhibition Enhance AMPH-Induced Locomotion and NVDR

*Fn* secretes multiple SCFAs, including butyrate ^45^. To confirm that our strain of *Fn* produced substantial levels of butyrate, we analyzed supernatants from growing cultures and found 3680 mg/L compared to 11 mg/L in unconditioned media (Fig. S2A). We next determined that oral administration of *Fn* increases overall fly butyrate content (Fig. S2B) and that oral administration of butyrate alone increased butyrate levels in fly heads (Fig. S2C). Considering that butyrate is a potent HDAC inhibitor^46^ and that HDAC inhibition potentiates AMPH-induced behavioral sensitization^27^, we hypothesized that butyrate mediates the ability of *Fn* to enhance AMPH behavioral properties and NVDR. In support of this hypothesis, we show that gnotobiotic flies orally treated with butyrate (25 mM) for 48 hours display both restored hyperlocomotion in response to AMPH (Fig. 2A) and increased NVDR (Fig. 2B). Furthermore, oral treatment of gnotobiotic flies (48 hours) with Trichostatin A (TSA, 1 µM), another HDAC inhibitor, closely paralleled the effects of butyrate (Fig. 2C, D), further suggesting that butyrate enhances these AMPH properties by HDAC inhibition. In contrast, oral administration of either acetate or propionate, additional SCFAs produced by *Fn*^45^ did not restore AMPH-induced hyperlocomotion and NVDR in gnotobiotic flies (Fig. S3A-D). These data suggest that *Fn* regulates AMPH actions *via* inhibition of HDACs, specifically through butyrate secretion. However, it is possible that butyrate might enhance NVDR through acute activation of a cell membrane receptor, in a manner analogous to fatty acid activation of free fatty acid receptor 2 and 3 in mammals^58^. To test for this possibility, we exposed isolated fly brains to an external solution containing either vehicle or butyrate (1 mM) for 10 min and then recorded AMPH-induced NVDR. Acute butyrate treatment did not enhance NVDR (Fig. S3E), suggesting that cell membrane receptors are not involved in butyrate regulation of this AMPH property.

**Figure 2:**
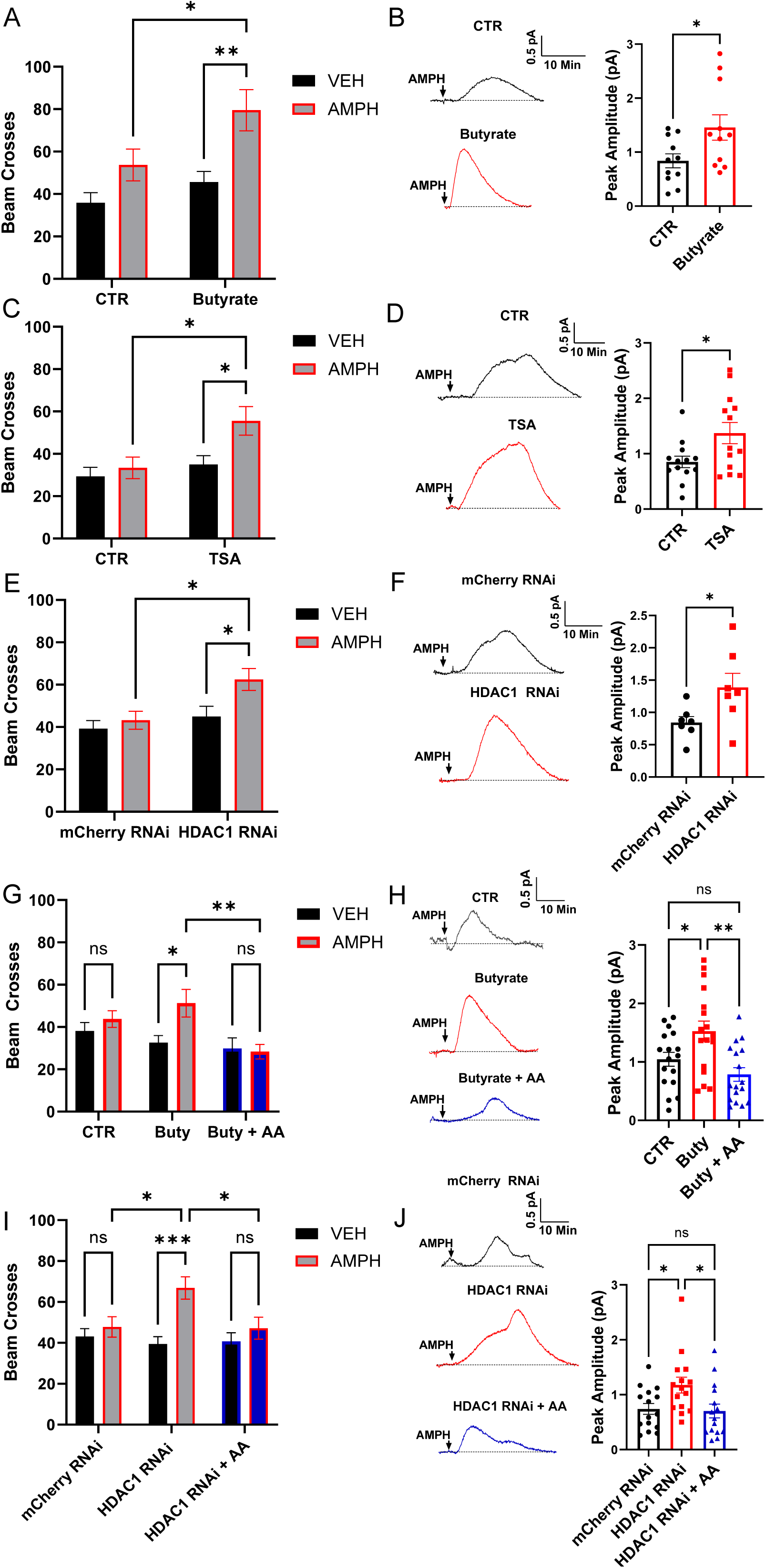
HDAC inhibition enhances AMPH responses. *Drosophila* were treated with either abx (CTR), abx with a pharmacological agent (panels A, B, C, D, G, H), or abx combined with RNAi with/out a pharmacological agent (panels E, F, I, J). Locomotor activity (left column) was measured as in Fig. 1 upon oral treatment of either VEH (black bars) or AMPH (10 mM, grey and blue bars). AMPH (20 µM)-induced NVDR (right column) was determined and quantified as in Fig. 1. **A.** Oral administration of butyrate (25 mM) significantly increased AMPH- induced hyperlocomotion (F_1, 182_ = 6.570 for effect of butyrate, p = 0.0112; F_1, 182_ = 13.89 for effect of AMPH, p = 0.0003, n = 45-47). **B.** Butyrate significantly augmented AMPH-induced NVDR (*t* = 2.306, p = 0.0320, n = 11). **C.** AMPH-induced locomotion is significantly increased by oral administration of TSA (F_1, 179_ = 7.164 for effect of TSA, p = 0.0081; F_1, 179_ = 5.607 for effect of AMPH, p = 0.0190, n = 45-47). **D.** TSA significantly enhanced AMPH- induced NVDR (*t* = 2.393, p = 0.0249, n = 13). **E.** HDAC1 RNAi (BDSC #31616) expressing flies display increased AMPH-induced hyperlocomotion with respect to mCherry RNAi (control) flies (F_1, 180_ = 5.683 for effect of AMPH, p = 0.0182; F_1, 180_ = 7.648 for effect of genotype, p = 0.0063, n = 43 – 48). **F.** AMPH-induced NVDR is significantly enhanced by RNAi targeting HDAC1 (*t* = 2.288, p = 0.0410, n = 7). **G.** AA significantly blunted the ability of butyrate to augment AMPH-induced hyperlocomotion (F_2, 361_ = 4.997 for effect of butyrate/AA; p = 0.0072; F_1, 361_ = 4.171 for effect of AMPH, p = 0.0419, n = 59-61). **H.** Butyrate significantly increased AMPH- induced NVDR and this increase was blocked by co-administration of AA (F_2, 54_ = 7.793, p = 0.0011, n = 19). **I.** AA significantly blunted HDAC1 RNAi ability to enhance AMPH-induced hyperlocomotion (F_2, 447_ = 3.743 for effect of interaction, p = 0.0244; F_1, 447_ = 11.42 for effect of AMPH, p = 0.0008, n = 73-78). **J.** HDAC1 RNAi expressing flies have significantly increased NVDR and this increase was blocked by AA (F_2, 42_ = 4.510, p = 0.0168, n = 15). Data is presented as mean ± SEM. Two-way ANOVA with Tukey’s multiple comparison test (A, C, E, G, I); Student’s unpaired t-test (B, D, F), One-way ANOVA with Tukey’s multiple comparison test (H, J).

To identify the HDAC isoform which inhibition drives the increase in AMPH response, we utilized the UAS/TH- GAL4 system to express RNAi constructs targeting the different *Drosophila* HDAC isoforms, specifically in DA neurons. As a control, we utilized an UAS-mCherry RNAi line. Among the different HDACs RNAi utilized, only the UAS-HDAC1 RNAi led to both an increase in AMPH-induced hyperlocomotion (Fig. 2E) as well as enhanced NVDR (Fig. 2F). Indeed, RNAi against the other HDACs isoforms as well as Sirtuin 2 (Sirt2), an NAD-dependent histone deacetylase, failed to recapitulate enhancement of these AMPH responses (Fig. S4A-H). Furthermore, we validated the role played by HDAC1 in NVDR with a second HDAC1 RNAi line (BDSC #34846; mCherry RNAi = 0.586±0.097, HDAC1 RNAi = 1.067±0.146; *t* = 2.742, *p* = 0.016, n = 9 by Student’s unpaired t-test). To verify whether butyrate and HDAC1 RNAi might enhance AMPH-induced hyperlocomotion and NVDR by increasing H acetylation, we tested whether blockage of acetylation, by anacardic acid (AA), alters the ability of HDAC inhibition to regulate these AMPH actions. AA is a histone acetyltransferase (HAT) inhibitor which potently reduces histone 3 (H3) acetylation in *Drosophila*^59^. We found that oral administration of AA in conjunction either with butyrate treatment (Fig. 2G-H) or to HDAC1 RNAi (Fig. 2I-J), significantly reduced the ability of these treatments to enhance AMPH psychomotor properties and NVDR.

### *Fn* colonization and Butyrate Exposure Increase DAT Expression

In cells, butyrate and other HDAC inhibitors enhance DAT expression^48,49,60^ by increasing acetylation of K9 and K14 of H3 within the DAT promotor^49^. As DAT expression positively correlates with the ability of AMPH to promote its psychomotor actions and NVDR^31,50^, and HDAC inhibition supports AMPH actions (Fig. 2), we first analyzed RNA sequencing data, from the Zhou *et al.* study^61^, and compared sterile flies treated with either vehicle or butyrate. Interestingly, butyrate treatment shifted the overall transcriptional profile of sterile *Drosophila* (Fig. 3A), upregulating the expression of 3732 genes and down regulating the expression of 3850 genes (Fig. 3B). Among the genes with a significant change in expression, several are involved in regulating DA homeostasis, including an elevation of dDAT expression (Fig. 3C). These findings led us to examine dDAT expression levels in heads isolated from gnotobiotic flies fed either vehicle, *Fn*, or butyrate. We found that *Fn* and butyrate feeding caused an increase in dDAT mRNA expression with respect to vehicle feeding (Fig. 3D-E). *Via* western blot analysis, we also found that both *Fn* (Fig. 3F) and butyrate (Fig. 3G) administration increases central dDAT protein expression. In order to describe changes in dDAT mRNA expression induced by either *Fn* or butyrate treatment, within an anatomical context, we performed RNAscope targeting dDAT. dDAT mRNA transcripts were identified based on thresholded-integrated density signals (see methods) and quantified across the entire brain within the ROIs selected, which include the posterior antennal lobe (PAL) and protocerebral anterior medial (PAM) neurons (Fig. 3H-J, circles) and normalized to the average of respective controls. Representative brain images of flies treated either with vehicle (Fig. 3H), inoculated with *Fn* (Fig. 3I), or fed butyrate (Fig. 3J). Both *Fn* colonization (Fig. 3K) and butyrate treatment (Fig. 3L) increases dDAT mRNA brain transcripts. To begin to determine whether *Fn* regulation of DAT expression is phylogenetically conserved, we utilized a *C. elegans* strain containing a T2A ribosome skip sequence followed by mNeonGreen, inserted downstream of the DAT coding region within the endogenous locus. Three days oral administration of *Fn* increased NeonGreen expression in cephalic sensilla (CEP) DA neurons (Fig. S5A-B), suggesting that *Fn* regulation of DAT expression is conserved across multiple host species.

**Figure 3:**
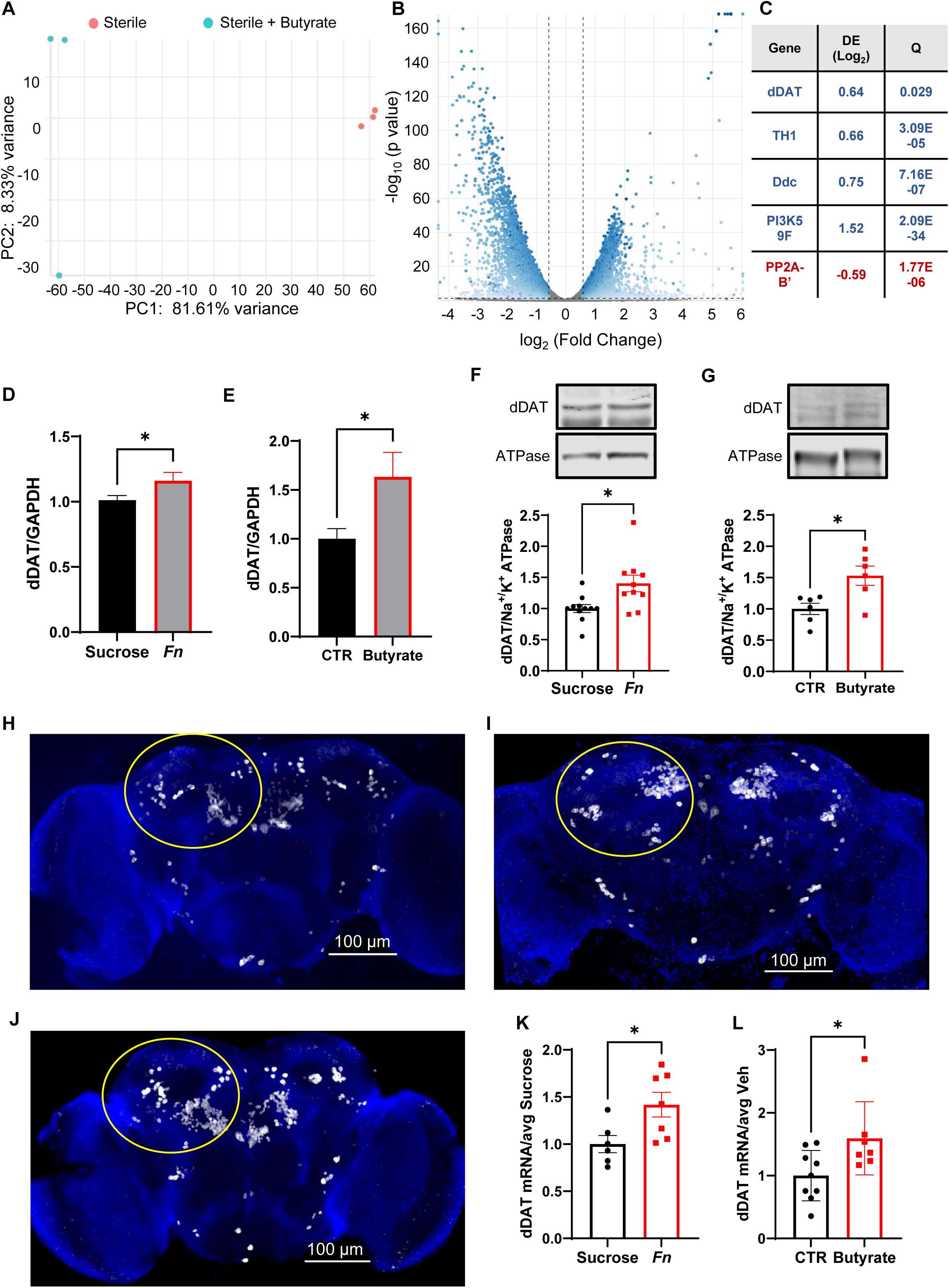
*Fn* colonization and butyrate treatment increase dDAT expression. **A-C.** Analysis of gene expression in sterile *Drosophila* with and without butyrate treatment; Principal component analysis (**A**) volcano plot (**B**), and table representing significantly increased (blue) and significantly decreased (red) of expression of genes within the DA network (**C**). **D.** dDAT mRNA expression levels normalized to GAPDH mRNA obtained from heads of gnotobiotic flies fed either sucrose or sucrose + *Fn* (*t* = 2.098, p = 0.0430, n = 17-19) **E.** dDAT mRNA levels normalized to GAPDH mRNA from heads of gnotobiotic flies treated orally with either vehicle (CTR) or butyrate (25 mM) (*t* = 2.339, p = 0.0267, n = 15). **F.** Representative immunoblots of dDAT and Na^+^/K^+^ ATPase (top) and their quantitative analysis (bottom) from heads of flies fed either sucrose (CTR) or inoculated with *Fn.* Data are normalized to the corresponding Na^+^/K^+^ ATPase level (*t* = 2.825, p = 0.0108, n = 10-11). **G.** Representative immunoblots of dDAT and Na^+^/K^+^ ATPase (top) and quantitative analysis (bottom) from brain of flies treated with either vehicle or butyrate (25mM). Data are normalized as in F (*t* = 2.952, p = 0.0145, n = 6). **H-I-J.** Representative RNAscope brain images of flies either fed sucrose, inoculate with *Fn*, or fed butyrate, respectively. **K.** Quantitation of brain dDAT mRNA *via* RNAscope in flies treated either with *Fn* or sucrose (*t* = 2.544, p = 0.0273, n = 6-7). **L.** Quantitation of brain dDAT mRNA (*via* RNAscope) in flies treated either with vehicle (CTR) or butyrate (t = 2.415, p = 0.0300, n = 7-9). Data is presented as mean ± SEM. Student’s unpaired t-test (D, E, F, G, K, L).

As previously shown in flies, knock out of dDAT (Fumin) eliminates the hyperlocomotive response to AMPH^34^ (Fig. S6A), consistent with what is observed in mice^62,63^. Furthermore, in fumin flies neither *Fn n*or butyrate rescue AMPH-induced hyperlocomotion (Fig. S6A-B), underscoring that changes in dDAT expression are pivotal for both *Fn* and butyrate regulation of AMPH-induced hyperlocomotion. *Fn* and butyrate regulation of AMPH- induced hyperlocomotion is psychostimulant specific as both treatments have no effect on responses to cocaine (COC) exposure (Fig. S7A-B). Indeed, COC functions as a DAT blocker and, therefore, an increase in DAT expression should not enhance the actions of COC^31,50^. Our data also demonstrate that COC-induced hyperlocomotion is insensitive to Abx treatment (Fig. S7B) possibly restricting changes in the fly microbiota to specific AMPH actions.

It is possible that our observed behavioral/electrochemical data could not only be due to increased dDAT expression, as we hypothesize, but that a general increase in brain DA content could also contribute to the enhanced AMPH behavioral phenotypes. Using mass spectrometry, we found that neither abx nor butyrate had any effects on the levels of biogenic amines (Fig. S8). Furthermore, it is possible that butyrate increases AMPH availability in the brain by altering the blood brain barrier (BBB) permeability^64^. However, butyrate treatment had no effect on concentration of AMPH found in *Drosophila* heads after AMPH treatment (Fig. S8).

### *Fn* and Butyrate Enhance AMPH-induced Courtship Behavior

AMPH, which stimulates DA neurotransmission, is commonly used to increase sexual motivation and desire in humans^65–67^. Consistently, in *Drosophila*, male courting behavior is also regulated by DA^68–71^ as well as DAT function^72^. Indeed, we have recently shown that treatment of naïve *Drosophila* males with AMPH enhances courtship^72^ and that, this enhancement is associated to the ability of DAT to support an increase in NVDR^72^. Consistent with the concept that NVDR regulates sexual motivation^72^, Abx treatment significantly decreased the ability of AMPH to induce courting (Fig. 4A; Svideo 1-2). Importantly, colonization of gnotobiotic *Drosophila* males with *Fn* or their oral treatment with butyrate restored the ability of AMPH to promote courting to levels similar to those observed in untreated flies (Fig. 4B-C; Svideo 3-6).

**Figure 4:**
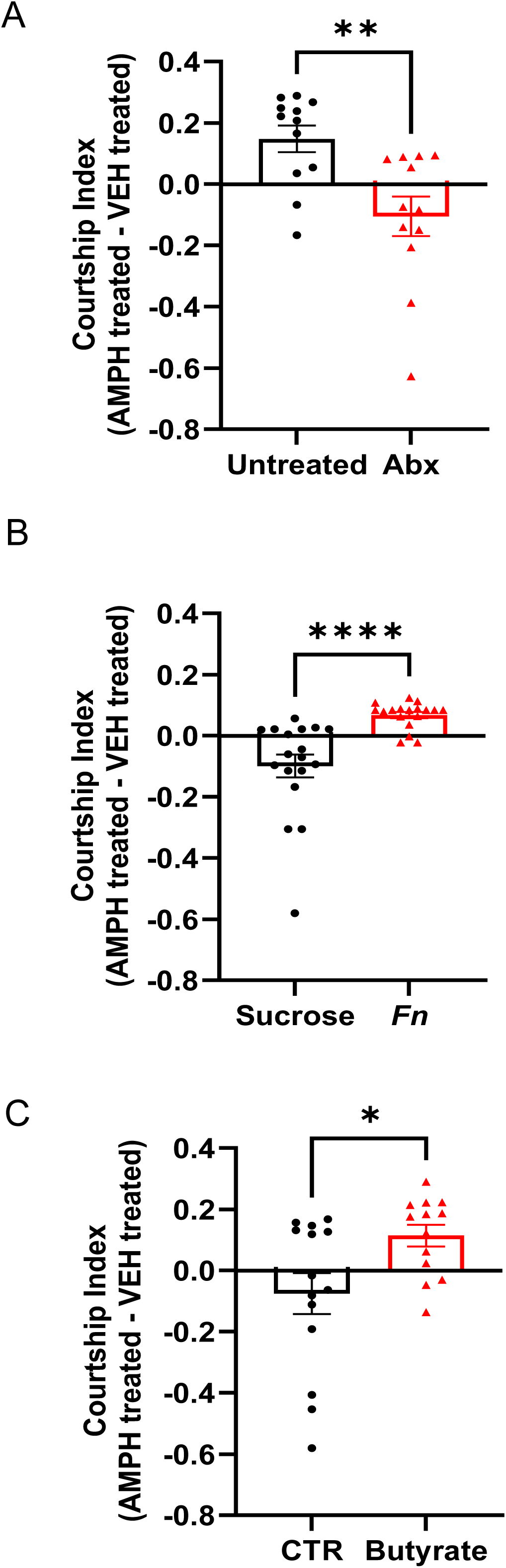
Both *Fn* inoculation and butyrate treatment enhance AMPH-induced courtship behavior. **A.** *Drosophila* males were either untreated or treated with abx. Male were then fed either vehicle or AMPH (10 mM) for 20 minutes prior to being introduced an unmated female fly. The courtship index (see methods) determined under vehicle treatment was subtracted from the index obtained upon AMPH exposure for both untreated (black bar) and abx treated (red bar). Abx treatment significantly reduced AMPH-induced courtship (U = 24, p = 0.0045, n = 12). **B.** Gnotobiotic males were fed either sucrose or colonized with *Fn*. After inoculation, flies were treated with either vehicle or AMPH and the courtship index determined, as in A. *Fn* inoculation (red bar) significantly increased AMPH-induced courtship with respect to sucrose (black bar). (U = 22, p = 0 < 0.0001, n = 18). **C.** Gnotobiotic *Drosophila* males were fed either fly food (CTR) or fly food + butyrate (25mM). Male flies were then treated with either vehicle or AMPH (10 mM) and the courtship index analyzed as in A. Butyrate treatment (red bar) significantly increased AMPH-induced courtship with respect to CTR treatment (black bar) (U = 41, p = 0.0145, n = 13-14). Data is presented as mean ± SEM. Mann-Whitney Test (A-C).

### Butyrate as well as HDAC1 inhibition Enhance AMPH Preference

Reward is regulated by DA across species^73^. Preference is a behavior that facilitates the transition from drug exposure to drug dependence and is used to determine the “rewarding” properties of a drug^74^. In *Drosophila*, we have shown that NVDR is required for AMPH to promote preference^32,72^. To measure preference, we used a two-choice capillary feeding apparatus (Fig. 5, left panels) and asked whether changes in microbiome composition regulates AMPH preference in *Drosophila* (see methods). After a day of acclimatation (Day 1), the baseline consumption of sucrose is recorded on day 2 in both the clear and blue capillary. On day 3 of the assay, AMPH is added to the blue capillary and preference for the blue capillary food is measured (Fig. 5A, left panels). Untreated flies demonstrated a robust preference for 250µM AMPH (increased consumption of food from the blue capillary normalized to total consumption) while gnotobiotic flies (Abx) had no preference (Fig. 5A, right). Interestingly, treatment of gnotobiotic flies with butyrate rescued AMPH preference with respect to vehicle (CTR) treated flies (Fig. 5B). Consistent with the notion that HDAC1 inhibition is involved in butyrate regulations of AMPH associated behaviors, expression of HDAC1 RNAi in DA neurons of gnotobiotic flies also rescued AMPH preference with respect to flies expressing mCherry RNAi (Fig. 5C). In the absence of AMPH, neither vehicle, butyrate, or HDAC1 RNAi flies display a demonstrable preference between the clear and blue capillary (Fig. S9A-C).

**Figure 5:**
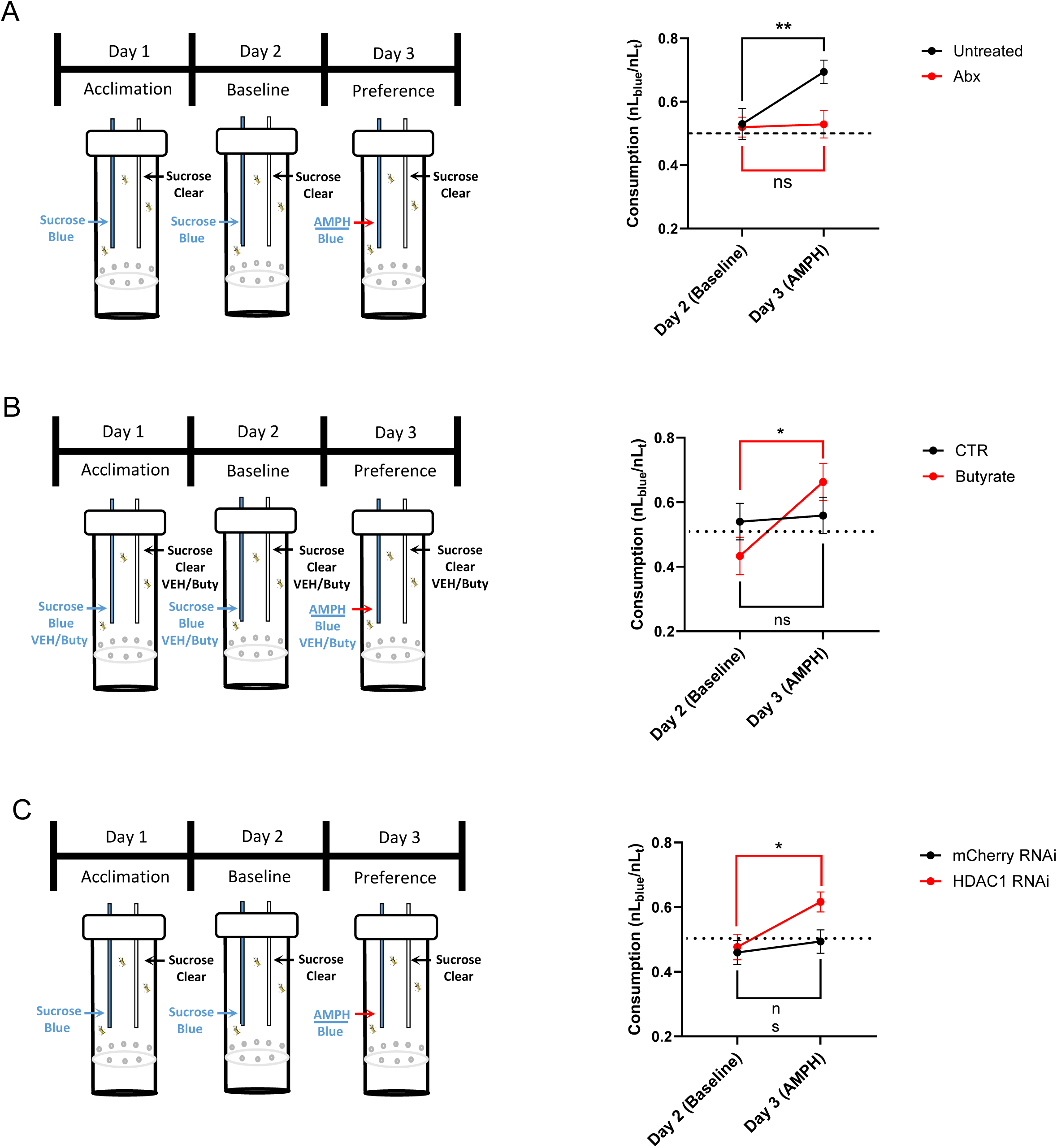
HDAC1 inhibition enhances AMPH preference. *Left column*: Cartoons illustrating the two-choice drug consumption assay for measuring AMPH preference in flies. The assay is comprised of three 24-hour testing periods: acclimation (Day 1), baseline (Day 2), and preference (Day 3). For the duration of the experiment, in half of the apparatuses, the capillaries contained sucrose in conjunction with a treatment condition (*e.g.* butyrate (Buty) or vehicle (CTR)). In each apparatus, one of the two capillaries has a blue dye in the food. On day 2 (Baseline), food consumption was recorded and normalized to total food consumed (blue capillary/blue+clear capillaries (nLblue/nLt). On day 3, AMPH (250µM) was added to the blue food capillaries. Food consumption was recorded again. Unless specified, both capillaries contain abx for the duration of the experiment. *Right column*: **A.** Untreated and abx treated flies (Abx; see methods) were given the choice of AMPH (blue capillary) or sucrose (clear capillary) on day 3. Untreated flies demonstrated a significant preference for AMPH as compared to abx treated flies. (F_1, 34_ = 5.519 for effect of day, p = 0.0248; F_1, 34_ = 4.469 for interaction, p = 0.0419, n = 18). **B.** Gnotobiotic flies were fed in both capillaries either vehicle (CTR) or butyrate (10mM) and given the choice of AMPH (blue capillary) on day 3. In contrast to vehicle treated (CTR), gnotobiotic flies treated with butyrate demonstrated a significant preference for AMPH on day 3. (F_1, 44_ = 5.109 for effect of day, p = 0.0288, n = 23). **C.** Gnotobiotic flies expressing either mCherry RNAi or HDAC1 RNAi were given the choice of AMPH on day 3. Only HDAC1 RNAi expressing flies demonstrated a significant preference for AMPH on day 3. (F_1, 43_ = 6.764 for effect of day, p = 0.0127, n = 22-23). Data is presented as mean ± SEM. Two-way Repeated Measures ANOVA with Šidák’s multiple comparisons test (A-C).

## Discussion

Altered gut microbiomes have been implicated in both changes in DA neurotransmission^2^ and substance use disorders^4^. Furthermore, several studies have associated exposure to psychostimulants with changes in gut microbiome composition^14–16,75^. In this context, it is important to note that antibiotic treatment regulates psychostimulant-induced behaviors^5,8,10^. Together, these data highlight the opportunity to utilize varying classes of antibiotics as well as probiotic interventions to therapeutically target PUD. However, this possibility is hindered by the absence of verified mechanisms of how specific bacterial species contribute to the expression of PUD and other brain disorders^53^. To begin to fill this knowledge gap is a recent study that points to COC-induced changes in the intestinal abundance of γ-Proteobacteria as a key regulator for the expression of COC behavioral responses^76^. In this study, the changes in γ-Proteobacteria abundance were linked to changes in intestinal glycine levels, highlighting a possible mechanism by which the microbiota might control COC-induced transcriptional plasticity^76^.

Using *Drosophila* as an animal model, we determined mechanistically how changes in the gut microbiota contribute to the expression of AUD. We identify *Fn,* a bacterium whose abundance is augmented by AMPH exposure^15,16^, as a key enhancer of AMPH-induced NVDR and associated behaviors, including sexual motivation and preference. We found that these *Fn* properties are likely mediated by *Fn* secretion of the SCFA butyrate. Consistently, VNP20009, a butyrate consumer^77^, does not regulate expression of AMPH-associated behaviors, further supporting the role of butyrate as the active molecule secreted by *Fn*. A major biological function of butyrate is HDAC inhibition and, consequently, modulation of gene expression. We found that specific inhibition of HDAC1 enhances the ability of AMPH to promote both NVDR and associated behaviors, a phenomenon we did not observe by selective knockdown of other HDAC isoforms. These data raise the possibility that *Fn* and butyrate selectively inhibit HDAC1 in order to regulate the expression of AMPH behaviors. We also demonstrate that both butyrate and *Fn* exposures increase dDAT expression and thereby NVDR in response to AMPH. Importantly, the psychomotor behavioral effects of *Fn,* butyrate, and HDAC1 inhibition are specific to the psychostimulant AMPH, since they do not affect the responses to COC, further supporting increases in dDAT expression as the mechanism driving *Fn* actions. Indeed, in *fumin* flies, both Fn and butyrate treatment failed in enhancing the psychomotor properties of AMPH. Finally, we show that *Fn*-mediated enhancement of DAT expression can occur across invertebrate species. Thus, we propose a feed-forward mechanism underlying the expression of AMPH behaviors, where AMPH exposure increases *Fn* abundance, which in turn promotes AMPH actions via synthesis of butyrate, HDAC1 inhibition, and upregulation of DAT expression. Based on these findings, while speculative, it is interesting to consider the possibility of *Fn* targeting therapies as treatment for AUD such as utilizing a phage specific to *Fn*^78^ to individually knock down this bacterium as a more focused treatment for AUD.

## Methods

### Bacteria Growth

*Fusobacterium nucleatum* (*Fn,* ATCC 23726) were grown in tryptic soy broth (TSB) (Millipore, Burlington, MA) supplemented with 1% Bacto Proteose Peptone No.3 (Gibco, Grand Island, NY) plus 0.05% freshly made cysteine hydrochloride (Sigma, Burlington, MA) (TSPC) in an anaerobic environment (Oxoid AnaeroJar 2.5L, Fisher, Waltham, MA) for 24-48 h under static conditions. *Fn* liquid cultures were grown for 48 hours before use. *E. coli* (OP50) and VNP20009 were grown in Luria-Bertani broth (LB) (RPI, Mt. Prospect, IL) shaking at 250 rpm at 37 °C for 16-24 hours.

### *Drosophila* Rearing and Stocks

All *Drosophila melanogaster* lines were maintained on standard cornmeal-molasses fly food (Nutrifly) at 25°C under a 12:12 h light-dark schedule. Fly stocks include Canton S (Bloomington Drosophila Stock Center (BDSC) #64349), TH-GAL4 (BDSC #8848), UAS-mCherry RNAi (BDSC #35785), UAS-HDAC1 RNAi (BDSC# 31616), UAS-HDAC1 RNAi (BDSC #34846), UAS-HDAC3 RNAi (BDSC #34778), UAS-HDAC4 RNAi (BDSC #34774), UAS-HDAC6 RNAi (BDSC #34072), UAS-Sirt2 RNAi (36868).

UAS-DAT RNAi (BDSC #31256), and *DAT^fmn^* (dDAT KO)^79^. For all crosses involving the UAS/GAL4 flies, female TH-GAL4 female flies were crossed with UAS males. Unmated adult male flies (3-9 days post eclosion) were used for all behavior and electrophysiological experiments (unmated adult female flies also used during the courtship assay; see below). ***C. elegans***: NFB2528 (dat-1(syb4741[dat-1::T2A::NeonGreen]) III; him-8(e1489) IV) *C. elegans* strain was used and cultured under standard conditions^80^.

### Gnotobiotic *Drosophila* generation

Unmated adult male flies (1-4 days post eclosion) were collected, anesthetized with CO_2_, and transferred to tubes containing standard fly food with an antibiotic cocktail (50µg/mL of ampicillin, erythromycin, and vancomycin) for 48 hours. After treatment with antibiotics, flies were anesthetized with CO_2_ and transferred to vials with ports for capillary feeding.

*Fn* Gut Inoculation: *Fn* was grown as a liquid culture (see above) at 7.0x10^5^ - 1.0x10^6^ cfu/µL. After growth of the bacteria, 2mL of the liquid culture was spun down (5,000rpm for 2 minutes) and the supernatant was removed. The pellet was resuspended in 200µL of sterile-filtered sucrose, which was added to the capillaries for feeding to the flies. Flies were fed 4 times a day (to refresh with healthy bacteria as *Fn* are anaerobic) for 72 hours.

VNP20009 Gut Inoculation: VNP20009 was grown as a liquid culture (see above) at 9.0x10^6^ - 1.1x10^7^ cfu/µL. Liquid culture of the bacteria was taken, spun down and prepared as above for *Fn*. Flies were fed once a day in the morning for 72 hours.

For all heat inactivation (HI) experiments, *Fn* was grown as above. 2mL of liquid culture was spun down (2,350 g for 2 minutes) and resuspended in 200µL of phosphate buffered saline (PBS). Bacteria were autoclaved for 45 minutes, spun down, PBS removed, and the pellets were resuspended in 200µL of sucrose, which was added to the capillaries for feeding flies. Flies were fed HI *Fn* once a day in the morning for 72 hours.

Colonization confirmation: After inoculation of bacteria for 72 hours, flies were externally sterilized by dipping in 100% EtOH, rinsed in PBS, and homogenized in PBS. Serial dilutions of the homogenates were plated on media plates or blood agar plates for determining cfus. Bacterial morphologies were observed under a microscope. Gram stain was used to verify typical morphology and characteristics of recovered *Fn* and confocal microscopy was used to verify the identity of VNP20009 in guts of Carnoy-fixed paraffin-embedded sections of flies using Hoechst 33342 (Invitrogen H3570), anti-*Salmonella* as the primary antibody (Abcam ab35156), and anti-rabbit- 488 (Invitrogen A11034) as the secondary antibody.

### Exposing C. elegans to Fn

Animals were egg-prepped according to the standard protocol^80^ and synchronized at the L1 stage, then allowed to grow to gravid adults on OP50 seeded NGM plates at 20°C. Gravid adults were then plated on NGM plates that were seeded with control or experimental bacteria. For seeding, bacteria were suspended in fresh LB broth to OD_600_ = 0.4. Control conditions contained only OP50. Experimental conditions contained 50% OP50 and 50% *Fn*. *Fn* were replaced 2-3 times per day (to refresh with healthy bacteria as *Fn* are anaerobic).

### Microscopy of *C. elegans*

Animals were immobilized using 150 mM sodium azide on 5% agarose. Imaging was performed on a spinning disk confocal microscope (Yokogawa W1 spinning disk mounted on a Nikon Eclipse Ti2) using the 20X air objective lens. Animals were imaged on day 1 after the first feeding and then again on day 3 after 6 total feedings. Maximum NeonGreen intensities of the CEP neuron cell bodies were quantified using Nikon imaging software.

### Drosophila Locomotion

Unmated adult male flies (1-4 days post eclosion) were collected, anesthetized with CO_2_, and transferred to vials containing standard fly food with either vehicle or an antibiotic cocktail (50µg/mL of ampicillin, erythromycin, and vancomycin, same as above) for 48 hours. In the case of experiments where the flies were treated with an additional compound (e.g. butyrate, trichostatin A (TSA), or anacardic acid (AA)), these were fed in conjunction with the antibiotics in the standard fly food. After this 48-hour period, flies were anesthetized with CO_2_ and transferred to glass tubes with standard fly food that contained the drug of interest (e.g. butyrate or TSA) as well as either 5mM COC or 10mM AMPH (or VEH). *Trikinetics Drosophila* Activity Monitoring (DAM) system (Waltham, Massachuesetts) was used to measure locomotion. After 1 hour of acclimation in the DAM system, locomotion was recorded for 100 minutes by beam breaks.

For locomotion post bacterial inoculation, unmated adult male flies (6-9 days post eclosion) who have been inoculated with bacteria (see above) were anesthetized with CO_2_ and transferred to glass tubes with standard fly food that contained either 5mM COC or 10mM AMPH (or VEH). After 1 hour of acclimation in the DAM system, locomotion was recorded for 100 minutes by beam breaks, as above.

### Drosophila Courtship

*Drosophila* courtship was measured in a custom 3D printed chamber (33x10x10mm). As previously described for the locomotion experiments, unmated adult male flies (1-4 days post eclosion) were collected, anesthetized with CO_2_, and transferred to vials containing standard fly food with either vehicle or an antibiotic cocktail (50µg/mL of ampicillin, erythromycin, and vancomycin, same as above) along with any additional compounds (e.g. butyrate) for 48 hours. After this 48-hour period, the adult male flies (3-6 days post eclosion) were starved for 6 hours in a vial with 1% agar. Flies were then fed standard fly food containing 10 mM AMPH (or VEH) for 20 minutes. Male flies were placed into the chamber for 10 minutes of acclimation prior to the introduction of an adult female fly (2-6 days post eclosion). Courtship was recorded for 10 minutes (Sony, ExmoreR OpticalSteadyShot). For courtship experiments done with flies inoculated with *Fn*, the protocol was the same as above but there was no 6-hour starvation period prior to the beginning of the experiment.

All courtship videos were scored blinded to the genotype and drug treatment. Video scorers manually timed and recorded the time spent displaying one of five courtship behaviors (following, wing extension, abdomen bend, tapping, and attempted copulation). The courtship index was calculated as the fraction of the amount of time the male spent courting the female over the 10 minutes. The normalized courtship was calculated by subtracting the average time spent courting after treatment with vehicle (standard fly food) from the time spent courting after AMPH treatment. Latency to court was measured as duration of time between introduction of the female and displaying of courtship behavior by the male. Achievement of copulation was recorded and utilized for analysis.

### *Drosophila* Two Choice AMPH Preference

A custom two choice apparatus was constructed in the lab to measure Drosophila drug preference^32^. Each apparatus contained two volumetric capillaries: one with clear food (100 mM sucrose) and one with blue food (100 mM sucrose, 500 µM blue dye). In these experiments, half of the flies also received vehicle, antibiotics, or butyrate in conjunction with the sucrose and dye (See Figure 6 for more details). Over a 72 hour period, consumption was measured and food was refreshed every 24 h. Adult male flies (3-6 days post eclosion) were anesthetized by CO_2_, transferred to the apparatus, and given 24 hours to acclimate to the liquid capillary food. The following 24 hours was used to calculate baseline preference of clear and blue food. On the third day, in order to determine drug preference, the blue capillary was supplemented with 250µM AMPH (See figure 6D-F). Preference was calculated as consumption of blue food over total consumption (blue capillary/blue + transparent capillaries; nLblue/nLt)

### *Drosophila* Brain Amperometry

Unmated adult male flies (1-4 days post eclosion) were collected and treated for 48 hours with antibiotics along with any additional compounds, as was described above for behavioral experiments. Brains from treated flies were dissected in cold Schneider’s *Drosophila* Medium (Gibco). Brains were then transferred to a sylgard dish (35mM) with 3mL of Lub’s external solution (130 mM NaCl, 10 mM HEPES, 34 mM dextrose, 1.5 mM CaCl_2_, 0.5 mM MgSO_4_, 1.3 mM KH_2_PO_4_, pH 7.35 and 300-310 mOsms/L) and pinned to the dish using tungsten write (California Fine Wire Company, Grover Beach, CA). A carbon fiber electrode (ProCFE; fiber diameter is 5 µm) (Axon Instruments, Australia) connected to an Axopatch 200B amplifier (Axon Instruments) was held at +600mV and placed between the PPL1 and PAM DA neuron regions. 20µM AMPH was used to elicit DA efflux. A sampling frequency of 100 Hz was used and the amperometric currents were low pass filtered at 10 Hz. Data were recorded and analyzed off-line using pCLAMP 9 software (Axon Instruments).

For experiments that involved acute butyrate treatment, the brain was pinned to the sylgard dish in the presence of Lub’s external solution (see recipe above) that was supplemented with 1mM sodium butyrate.

### RNA Sequencing analysis

RNA sequencing data were generated from previous studies^61^ that were deposited in the NCBI Gene Expression Omnibus database under accession number GSE169135. These data were analyzed to compare gene expression between sterile *Drosophila* treated with or without dietary supplementation with butyrate. Data were analyzed using the Basepair website and associated Differential Expression (DESeq2) platform with cutoffs set at adjusted *p*-value ≤ 0.05 and fold change ≥ 1.5.

### PCR

Unmated adult male flies (1-4 days post eclosion) were collected and treated for 48 hours with antibiotics along with any additional compounds, as was described above for behavioral and electrophysiological experiments. For PCR post bacterial inoculation, unmated adult male flies (6-9 days post eclosion) who had been inoculated with bacteria (see above). After treatment/inoculation, fly heads were removed and immediately flash frozen on dry ice. 30 heads were used per sample. Samples were stored at -80°C until RNA extraction was performed.

RNA was extracted using a RNeasy Mini Kit (Qiagen, Germantown, MD). Frozen heads were homogenized using a 1.5mL pestle (VWR, Radnor, PA) in 350µL RLT buffer and then all other steps were performed according to the manufacturer’s protocol. RNA was eluted in 30µL of water. Quantity and quality of the RNA were assessed using a Nanodrop spectrophotometer (Nanodrop Technologies, Wilmington, DE). RNA was then stored at -80°C until further processing. cDNA was produced using iScript Reverse Transcription Supermix for RT-qPCR (Biorad, Hercules, CA) in 20µL total volume using 500 ng total RNA, following the manufacturer’s protocol. cDNA was stored at -20°C until used.

qPCR was performed in duplicates on 100ng total cDNA using iQ SYBR Green Supermix (Bio-Rad, Hercules, CA) on QuantStudio 6 Flex PCR System (Thermofisher). Relative normalized transcript level was determined by delta-delta Ct method. *Gapdh* was used as the normalizing gene. *DAT* primers: 5’ –GTCATTATTCCCTGGTCGTTG- 3’ and 5’ –GATTTCCCATGGCATACAAGTC -3’. *Gapdh* Primers: 5’ –CCTGGCCAAGGTCATCAATG- 3’ and 5’ –ATGACCTTGCCCACAGCCTT- 3’.

### RNAscope and Imaging

RNAscope processing and imaging was used to quantify transcript expression levels in PFA-fixed brains. Probes for RNAscope were designed by Advanced Cell Diagnostics (Newark, CA) (Table 1). All brains were processed using the RNAscope Assay V2 kit (Advanced Cell Diagnostics). Brains were dissected in ice cold Schneider’s Insect Media and processed according to published protocol^81^ with minor adaptations. Brains were fixed in 2% paraformaldehyde (PFA) in PBS at room temperature (RT) for 55 minutes under gentle agitation, washed in 0.5% Triton X-100 in PBS (PBT), then dehydrated in a graded series using ethanol and stored at 4°C overnight. Brains were subsequently rehydrated in a graded ethanol series on ice and then incubated with pre-chilled 5% acetic acid (5 min) and washed with pre-chilled PBS and then PBT. Brains were incubated with 1% sodium borohydride (30 min, 4°C) and then washed with PBT. Multiplex fluorescent *in situ* hybridization via RNAscope was performed in accordance with the manufacturer’s protocols (Advanced Cell Diagnostics). Briefly, the samples were first treated with peroxide (10 min, RT) and protease treatment (30 min, RT) followed by probe hybridization (2 hours, 40°C) and then probe amplification (40°C).

**Table 1:** List of RNAscope probes from Advanced Cell Diagnostics.

| Probe | Cat. Number | Information |
| --- | --- | --- |
| RNAscope® Probe - Dm-DAT - *Drosophila melanogaster* dopamine transporter transcript variant A (DAT) mRNA | 868771 | Target probe |
| RNAscope® Negative Control Probe - DapB - *Bacillus subtilis* strain SMY methylglyoxal synthase (mgsA) gene, partial cds dihydrodipicolinate reductase (dapB) gene, complete cds and YpjD (ypjD) gene, partial cds | 310043 | Negative Control |
| melanogaster glyceraldehyde 3 phosphate dehydrogenase 1 transcript variant A (Gapdh1) mRNA | 470801 | Positive control |

Hybridized samples were incubated with 0.025 mg/mL Hoeschst 33342 (30 min, RT) prior to mounting with Prolong Diamond Antifade Mountant (Invitrogen, Waltham, MA).

Images were collected at UAB’s High Resolution Imaging Facility with a Nikon A1R HD Confocal Microscope. Images were preprocessed in Advanced NIS-Elements for brain orientation, condition blinding, and anatomical labelling. Quantification was performed using integrity density thresholding within global ROIs using Fiji/ImageJ software (National Institutes of Health, Bethesda, MD)^82^. Brains were thresholded for complete background subtraction that captured all positive signal as optimized during control trials. Positive pixels were summed as a representative of levels of target transcripts. Results were normalized to gnotobiotic control flies.

### Western Blots

Fly heads were removed and immediately flash frozen on dry ice. 10-20 heads were pooled per sample. Frozen heads were homogenized using a 1.5mL pestle (VWR) in lysis buffer (1% SDS, 50 mM NaF). Lysates were sonicated and centrifuged at 15,000 x g for 30 minutes. Supernatants were then separated by 10% SDS-PAGE gel, transferred to the polyvinylidene fluoride membrane (PVDF) (Millipore, Bedford, MA) and immunoblotted. Blocking was done with Intercept blocking buffer (Licor, Lincoln, NE). Primary antibodies used were Anti-*Drosophila* DAT (1:500, clone 15A11, made in house and provided by Dr. Eric Gouaux) and Anti-Na^+^/K^+^ ATPase (1:1000, a5, DSHB, Iowa City, IA). The secondary antibody used was goat anti-mouse IgG 800CW (1:10000, 926-32210, Licor). Controls were averaged within each experiment and each sample normalized to its respective control group.

### Mass Spectrometry

*Biogenic Amines.* Stock solutions of each analyte of interest (5ng/μL each) were made in DI water and stored at -80°C. To prepare internal standards, stock solutions were derivatized in a similar manner to samples using isotopically labeled benzoyl chloride (^13^C_6_-BZC) as follows: 200uL of the stock solution was mixed with 400uL each of 500mM NaCO_3_ (aq) and 2% ^13^C_6_-BZC in acetonitrile was added to the solution. After two minutes, the reaction was stopped by the addition of 400uL 20% acetonitrile in water containing 3% sulfuric acid. The solution was mixed well and stored in 10uL aliquots at -80°C. One aliquot was diluted 100x with 20% acetonitrile in water containing 3% sulfuric acid to make the working internal standard solution used in the sample analysis.

Fly heads were removed and immediately flash frozen on dry ice. 20 heads were pooled per sample. Fly heads were homogenized, using an ultrasonic dismembrator, in 100-750 µl of 0.1M TCA, which contained 10 mM sodium acetate, 0.1 mM EDTA, and 10.5 % methanol (pH 3.8). Ten microliters of homogenate were used for protein quantification. Samples were spun in a microcentrifuge at 10,000 g for 20 minutes at 4°C. The supernatant was removed for LC/MS analysis.

Analytes in tissue extract supernatant were quantified using liquid chromatography/mass spectrometry (LC/MS) following derivatization with benzoyl chloride (BZC). 5uL of supernatant was then mixed with 10uL each of 500mM NaCO_3_ (aq) and 2% BZC in acetonitrile in an LC/MS vial. After two minutes, the reaction was stopped by the addition of 10uL internal standard solution.

LC was performed on a 2.1 x 100 mm, 1.6 µm particle CORTECS Phenyl column (Waters Corporation, Milford, MA, USA) using a Waters Acquity UPLC. Mobile phase A was 0.1% aqueous formic acid and mobile phase B was acetonitrile with 0.1% formic acid. MS analysis was performed using a Waters Xevo TQ-XS triple quadrupole tandem mass spectrometer. The source temperature was 150°C, and the desolvation temperature was 400°C. The LC gradient is shown below.

Protein concentration was determined using the BCA Protein Assay Kit (Thermo Scientific, Waltham, MA USA) in a 96-well plate format. 10uL of tissue homogenate was mixed with 200 μl of mixed BCA reagent per manufacturer instructions. The plate was incubated at 23°C for two hours before absorbance was measured by plate reader (POLARstar, Omega, Bienne, Switzerland).

Amphetamine: Standard & sample preparation involved generating a 7-point standard curve using an authentic reference standard (source unknown) with concentrations ranging from 0.1 to 100 ng/ml in 0.1% Formic Acid (FA). Fruit fly heads (30 / sample) were pooled and homogenized in 100 µl 80% methanol. Protein precipitation was achieved by adding 800 µl acetonitrile (MeCN) with 1% FA, followed by vortexing and solid-phase extraction using Phenomenex Phree SPE cartridges. Extracts were dried under nitrogen gas and reconstituted in 0.1% FA. Dilutions, 1:10 and 1:100, were also prepared to ensure signal intensity fell within the standard curve range. HPLC-MS/MS analysis was performed using a Shimadzu Prominence LC system coupled to a Sciex API 4000 triple quadrupole mass spectrometer. Chromatographic separation was achieved on using 5 µl injection on a Phenomenex Kinetex Biphenyl (P/N 00B-4622-AN) column at 40°C using a gradient elution (10 – 95% B over 3 minutes) with mobile phases A) 0.1% FA and B) MeCN in 0.1% FA at a flowrate of 0.4 ml/min. The MS was operated in positive electrospray ionization mode, monitoring the transitions m/z 136 → 44 and m/z 136 → 91. Data processing was performed using Sciex MultiQuant 3.0.2 software, and standard curves were generated by linear regression with 1/x^2^ weighting.

### *Butyrate.* Extraction

Flies (n=20) or fly heads (n=20) were homogenized at 0°C by sonication (Fisher Sonic Dismembrator, 3 x 15 sec, power = 3) to a final tissue density of 50 mg/mL in PBS containing 10 % (v/v) MeOH. Insoluble debris was removed by centrifugation (3,000 x g, 10 minutes, 4°C); supernatants were then transferred to clean Eppendorf tubes and stored at -20°C until analysis.

### LC-HRMS Analysis

Butyric acid (BA) and a deuterium-labeled internal standard (BA-*d*_5_) were derivatized with the reagent dansyl hydrazine and the carboxyl activating agent 1-Ethyl-3-(3-dimethylaminopropyl)carbodiimide (EDC) to their corresponding dansyl hydrazide derivatives. Stock solutions of EDC were made up fresh in water and used immediately. Briefly, 100 mL aliquots of thawed extracts were spiked with 1.25 nmol BA-*d*_5_, diluted with 100 mL of water/acetonitrile (3:2) buffered with 250 mM sodium phosphate pH 2.5, and derivatized at room temperature with dansyl hydrazine (25 mL x 50 mg/mL) and EDC (25 mL x 150 mg/mL). After two hours at RT, 25 mL of 5 % (*v/v*) trifluoroacetic acid in water was added to quench the excess EDC. Following centrifugation (10,000 x *g*, 30 min, 5 °C), quenched reaction mixtures were transferred to 2-mL autosampler vials equipped with low-volume polypropylene inserts and Teflon-lined rubber septa for analysis. The sample injection volume was 10 mL. BA calibration standards were prepared in water and derivatized in the same manner. LC-MS analysis was performed using a Thermo Q Exactive HF hybrid quadrupole/orbitrap high resolution mass spectrometer interfaced to a Vanquish Horizon HPLC system (Thermo Scientific). The mass spectrometer was operated in negative ion mode, with targeted selected ion monitoring (t-SIM) detection of specified precursor ions at a resolving power of 60,000, an isolation window of 2.0 *m/z*, and the following APCI source parameters: corona discharge current 15 mA; ion transfer tube temperature 300°C; auxiliary gas temperature 100 °C; *S*-lens RF level 65; N_2_ sheath gas 35; N_2_ auxiliary gas 10; in-source CID 10 eV. Extracted ion chromatograms were constructed for BA and BA-*d*_5_ with the following exact masses and a mass tolerance of +/- 5 ppm: BA: [M-H]^-^ 334.1231; BA-*d*_5_: [M-H]^-^ 339.1545. Data acquisition and quantitative spectral analysis were done using Thermo Xcalibur version 4.1.31.9 and Thermo LCQuan version 2.7, respectively. Calibration curves were constructed by plotting peak areas against analyte concentrations for a series of eight calibration standards, ranging from 0.002 to 10 total nmol BA. A weighting factor of 1/C^2^ was applied in the linear least-squares regression analysis to maintain homogeneity of variance across the concentration range. An Acquity BEH C18 reverse phase analytical column (2.1 x 100 mm, 1.7 mm, Waters, Milford, MA) was used for all chromatographic separations. Mobile phases were made up of 15 mM ammonium acetate and 0.2 % acetic acid in (A) water/acetonitrile (9:1) and in (B) acetonitrile/methanol/water (90:5:5). Gradient conditions were as follows: 0–1.0 min, B = 0 %; 1–8 min, B = 0–100 %; 8–10 min, B = 100 %; 10–10.5 min, B = 100–0 %; 10.5–15 min, B = 0 %. The flow rate was maintained at 300 mL/min, and the total chromatographic run time was 15 min. A software-controlled divert valve was used to transfer the LC eluent from 0 to 5 min and from 8 to 15 min of each chromatographic cycle to waste.

### Gas Chromatography

Supernatants (2 mL) from bacterial cultures were freeze-dried (Labconco Freezone 2.5; Kansas City, MO, USA) approximately for 14 hours with collector temperature is set to 60 °C. Fatty acid methyl esters (FAMES) were obtained by trans-methylation of the freeze-dried samples as previously described^83^. FAMES were separated using gas chromatography (Agilent GC-FID 8890; Santa Clara, CA, USA) and identified by comparison with previously characterised standards and GC-MS (Agilent 5977C GC/MSD; Santa Clara, CA 95051).

### Statistics

Experiments were designed after statistical power calculations based on preliminary data using GPower 3.1. Statistical analyses were run on GraphPad Prism 9.1 (San Diego, CA). Shapiro-Wilk and Kolmogorov-Smirnov normality tests were performed to ensure data was normally distributed and Bartlett’s tests were performed to check for heteroscedasticity. If data failed these assumptions the proper non-parametric tests were utilized. Unpaired t-tests were used for all comparisons between two groups and Welch’s correction was used whenever the variances significantly differed between groups. When using one-way or two-way ANOVA’s, Tukey’s post hoc was used. For all repeated measures two-way ANOVA’s, Šidák’s multiple comparisons post hoc was used. For all experiments * = *p* < 0.05, ** = *p* < 0.01, *** = *p* < 0.001, **** = *p* < 0.0001.

## ACKNOWLEDGEMENTS

Research reported in this publication was supported by the UAB High Resolution Imaging Facility and the Targeted Metabolomics and Proteomics Laboratory (UAB Health Services Foundation General Endowment Fund).

## AUTHOR CONTRIBUTIONS

SJM, XC, HL, HS, HW, AMC, and AG conceptualized the study. SJM, XC, YZ, CR, ST, SP, CGA, TR, DPS, AE, SNL, CEH, and AMC performed the experiments and analyzed the data. All authors contributed to data interpretation and figure preparation. SJM, AMC, and AG wrote the manuscript. HL, HS, HW, AMC, and AG supervised the study.

## FUNDING

This work was supported by T32NS061788 (SJM), T32GM135028 (CR and CEH), a NIDA Diversity Supplement to R01DA038058 (AMC and AG), R01DA038058 (AG), R01DA056484 (AMC and AG), R21DA059844 (AMC), R00HD098371 (HS) and the HHMI Freeman Hrabowski Scholar program (HS).

**Supp. Fig. 1. Colonization of *Drosophila* with VNP20009 does not augment psychomotor responses to AMPH**. **A.** VNP20009 (VNP) fed flies were assessed for colonization by homogenization and plating; representative images of plating (*left-top*) and overall cfu/fly (*left-bottom*). Sucrose, Fn, and VNP fed flies were additionally assessed for VNP colonization by immunofluorescent imaging following staining with anti-*Salmonella* (*right*). **B.** VNP20009 colonization does not augment the ability of AMPH to cause hyperlocomotion (F_1, 139_ = 24.35 for effect of AMPH, p < 0.0001, n = 32-38). **C**. VNP colonization does not increase AMPH-induced NVDR (t = 0.9689, p = 0.3491, n = 7). Data is presented as mean ± SEM. Two-way ANOVA with Tukey’s multiple comparison test (B); Student’s unpaired t-test (C).

**Supp. Fig. 2. *Fn* produces butyrate and leads to elevated levels in flies. A.** GC analysis demonstrating butyrate is elevated in supernatants from cultures of *Fn* compared to cultures of VNP and unconditioned media (TSB). **B.** MS analysis demonstrating butyrate levels are elevated in gnotobiotic flies administered *Fn* (t = 2.960, p = 0.0253, n = 4). **C.** MS analysis demonstrating butyrate levels are elevated in heads of flies orally administered 25 mM butyrate (t = 2.428, p = 0.0414, n = 5). Data is presented as mean ± SEM. Student’s unpaired t-test (A, B).

**Supp. Fig. 3. Acetate and propionate do not enhance AMPH actions**. *Drosophila* were treated with either abx (CTR) or abx combined with a pharmacological agent. (A, C) Locomotor activity was measured as in Fig. 1 upon oral treatment of either VEH (black bars) or AMPH (10 mM, grey bars). (B, D, E) AMPH-induced NVDR was determined and quantified as in Fig. 1. **A.** There was no significant difference in AMPH-induced locomotion after oral acetate (50mM and 75mM) treatment (F_1, 259_ = 15.51 for effect of AMPH, p = 0.0001, n = 42 - 48). **B.** There was no significant difference in AMPH-induced NVDR after oral acetate treatment. (U = 49, p = 0.97105, n = 10). **C.** There was no significant difference in AMPH-induced locomotion after oral administration of propionate (25mM and 50mM) treatment (F_1, 253_ = 5.452 for effect of AMPH, p = 0.0203, n = 39 - 48). **D.** There was no significant difference in AMPH-induced NVDR after oral administration of propionate (*t* = 0.3524, p = 0.7282, n = 11). **E.** There was no significant difference in AMPH-induced NVDR after acute (10 minutes) butyrate treatment (1mM) (U = 41, p = 0.5288, n = 10). Data is presented as mean ± SEM. Two-way ANOVA with Tukey’s multiple comparisons test (A, C); Mann-Whitney Test (B, E); Student’s unpaired t-test (D).

**Supp. Fig. 4. Multiple HDACs do not regulate AMPH-induced locomotion and NVDR**. (A, C, E, G) Locomotor activity measured as in Fig. 1 upon oral treatment of either VEH (black bars) or AMPH (10 mM, grey bars) in gnotobiotic flies expressing either a specific HDACs or mCherry (control) RNAi. (B, D, F, H) In the same flies AMPH-induced NVDR was determined and quantified as in Fig. 1. **A.** HDAC3 RNAi expressing flies did not demonstrate increased AMPH-induced locomotion (F_1, 181_ = 5.769 for effect of AMPH, p = 0.0173; F_1, 181_ = 4.660 for effect of genotype, p = 0.0322, n = 42 – 48). **B.** HDAC3 RNAi expressing flies did not demonstrate increased AMPH-induced NVDR (*t* = 0.0255, p = 0.9801, n = 7). **C.** AMPH significantly increased locomotion in mCherry RNAi expressing flies but failed to increase locomotion in HDAC4 RNAi expressing flies (F_1, 180_ = 6.108 for effect of AMPH, p = 0.0144, n = 44 – 48). **D.** HDAC4 RNAi expressing flies did not demonstrate increased AMPH- induced NVDR (U = 22, p = 0.8048, n = 7). **E.** AMPH significantly increased locomotion in mCherry RNAi expressing flies compared to HDAC6 RNAi expressing flies (F_1, 179_ = 7.532 for effect of AMPH, p = 0.0067; F_1, 179_ = 11.21 for effect of genotype, p = 0.001, n = 43 – 48). **F.** HDAC6 RNAi expressing flies did not demonstrate increased AMPH-induced NVDR (U = 21, p = 0.7104, n = 7). **G.** Sirt2 RNAi expressing flies did not demonstrate increased AMPH-induced locomotion (F_1, 178_ = 4.200 for effect of AMPH, p = 0.0419; F_1, 178_ = 5.303 for effect of genotype, p = 0.0224, n = 40 – 48). **H.** Sirt2 RNAi expressing flies did not demonstrate increased AMPH-induced NVDR (U = 16, p = 0.3176, n = 7). Data is presented as mean ± SEM. Two-way ANOVA with Tukey’s multiple comparisons test (A, C, E, G). Student’s unpaired t-test (B), Mann-Whitney test (D, F, H).

Supp. Fig. 5**. *Fn* increases DAT-1 expression in *C. elegans* in DA CEF neuron cell bodies.** DAT-mNeonGreen reporter worms were fed either *Escherichia coli* (OP50) or OP50 + *Fn* for 3 days. Fluorescence intensity was measured on days 1 and 3. **A.** Representative images from day 3 with CEP neuron cell bodies denoted. **B.** Quantitated fluorescence in CEP neuron cell bodies, normalized to the average fluorescence of the control within each day. Data is presented as mean ± SEM. Two-way ANOVA with Tukey’s multiple comparisons test (F_1, 33_ = 9.025 for effect of *Fn*, p = 0.0051; F_1, 33_= 5.227 for effect of day, p = 0.0288, n = 8-10).

**Supp. Fig. 6. In Fumin flies *Fn* and butyrate fail in upregulating AMPH actions**. **A.** Fumin flies were administered either sucrose or *Fn*. Then, flies were treated with either VEH (black bars) or AMPH (10mM, grey bars) and locomotor activity measured by beam crosses. AMPH significantly decreased locomotion in the *Fn* group. (F_1, 80_ = 11.55 for effect of AMPH, p = 0.0011, n = 19 – 23). **B.** Fumin flies were fed either butyrate (25mM) or vehicle (CTR) as described above. After treatment flies were treated with vehicle (black bars) or AMPH (10mM, grey bars) and locomotor activity recorded. (F_1, 119_ = 0.2544 for effect of AMPH, p = 0.6149, n = 30 – 32). Data is presented as mean ± SEM. Two-way ANOVA with Tukey’s multiple comparisons test (A, B).

**Supp. Fig. 7. *Fn colonization* and butyrate treatment do not enhance COC-induced hyperlocomotion. A.** Flies were fed either sucrose or sucrose + *Fn* and treated either with vehicle (black bars) or COC (5 mM, grey bars) and locomotor activity recorded by beam crosses. COC significantly increased locomotion in both groups with no differences observed between COC groups. (F_1, 118_ = 60.65 for effect of COC, p < 0.0001, n = 29 – 32). **B.** Flies were either not treated (No Abx), treated with abx only (CTR), or treated with abx plus butyrate (Butyrate) and fed either vehicle (black bars) or COC (5mM, grey bars). Locomotion was recorded by beam crosses. COC significantly increased locomotion in all groups, however no differences were observed between COC groups (F_1, 313_ = 22.72 for effect of COC, p < 0.0001, n = 40 – 63). Data is presented as mean ± SEM. Two-way ANOVA with Tukey’s multiple comparisons test (A) or with Šidák’s multiple comparisons test (B).

**Supp. Fig. 8. Brain levels of either biogenic amines or AMPH after oral butyrate treatment**. **A.** Brain DA levels measured after vehicle or abx treatment (t = 0.8993, p = 0.3948, n = 5). **B.** DA levels measured in gnotobiotic flies after either vehicle (CTR) or butyrate (25mM) treatment (t = 0.2648, p = 0.7978, n = 5). **C.** 5-HT levels after either vehicle or abx treatment (t = 0.5412, p = 0.6031, n = 5). **D.** 5-HT levels in gnotobiotic flies after vehicle (CTR) or butyrate (25mM) treatment (t = 0.7491, p = 0.4923, n = 5). **E.** Brain octopamine (OA) levels detected after either vehicle or abx treatment (t = 0.3314, p = 0.7489, n = 5). **F.** In gnotobiotic flies butyrate (25mM) treatment doesn’t change brain OA levels with respect to vehicle (CTR) treatment (t = 0.9611, p = 0.3646, n = 5). **G.** No differences were detected in brain AMPH levels in flies fed AMPH and either vehicle (CTR) or butyrate (t = 0.4170, p = 0.6912, n = 4). Data is presented as mean ± SEM. Student’s unpaired t-test (A-C, E- F) with Welch’s correction (D, G)

**Supp. Fig. 9. Blue food is not associated with preference**. **A-C**. Preference for blue food in the absence of AMPH on day 3, in flies treated as in Fig. 5 using the two-choice capillary apparati. Unless specified, both capillaries contain abx for the duration of the experiment. **A.** Flies treated with either vehicle (untreated) or abx demonstrated no preference for either capillary (F_1, 32_ = 0.5778 for effect of day, p = 0.4527, n = 17). **B.** Flies treated with either vehicle (CTR) or butyrate demonstrated no preference for either capillary (F_1, 46_ = 0.0005 for effect of day, p = 0.9820, n = 24). **C.** Flies expressing either mCherry RNAi or HDAC1 RNAi demonstrated no preference for either capillary (F_1, 45_ = 0.1430 for effect of day, p = 0.7071, n = 23-24).

