## Supplementary figures and images for "*Fusobacterium nucleatum* determines the expression of amphetamine-induced behavioral responses through an epigenetic phenomenon"

### Supplemental Figure 1

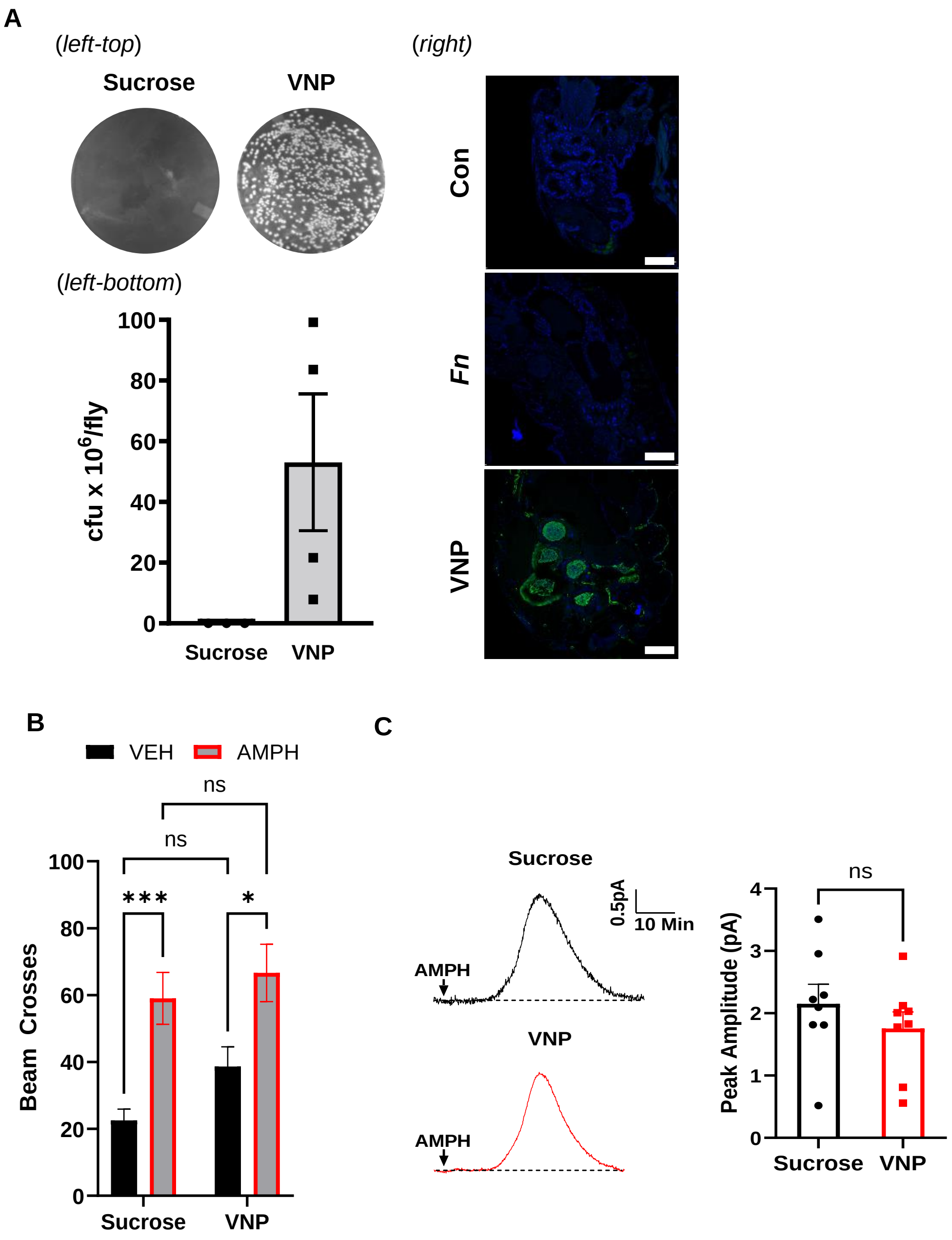

### Supplemental Figure 2

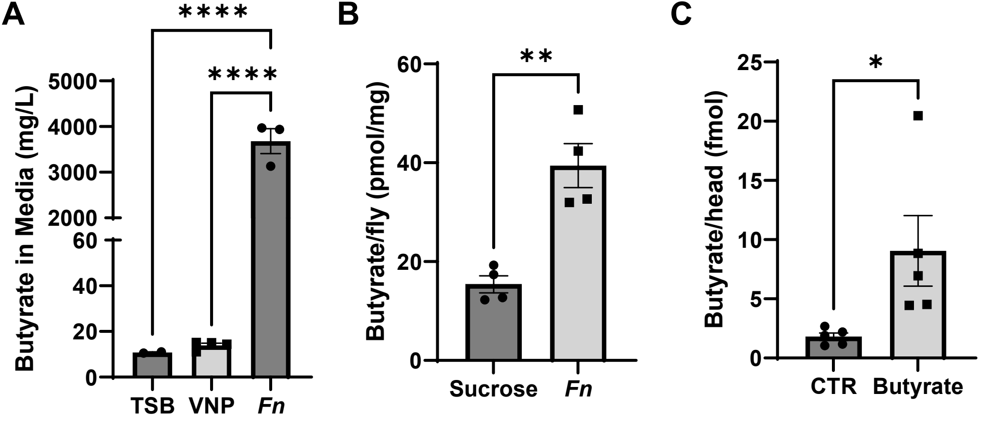

### Supplemental Figure 3

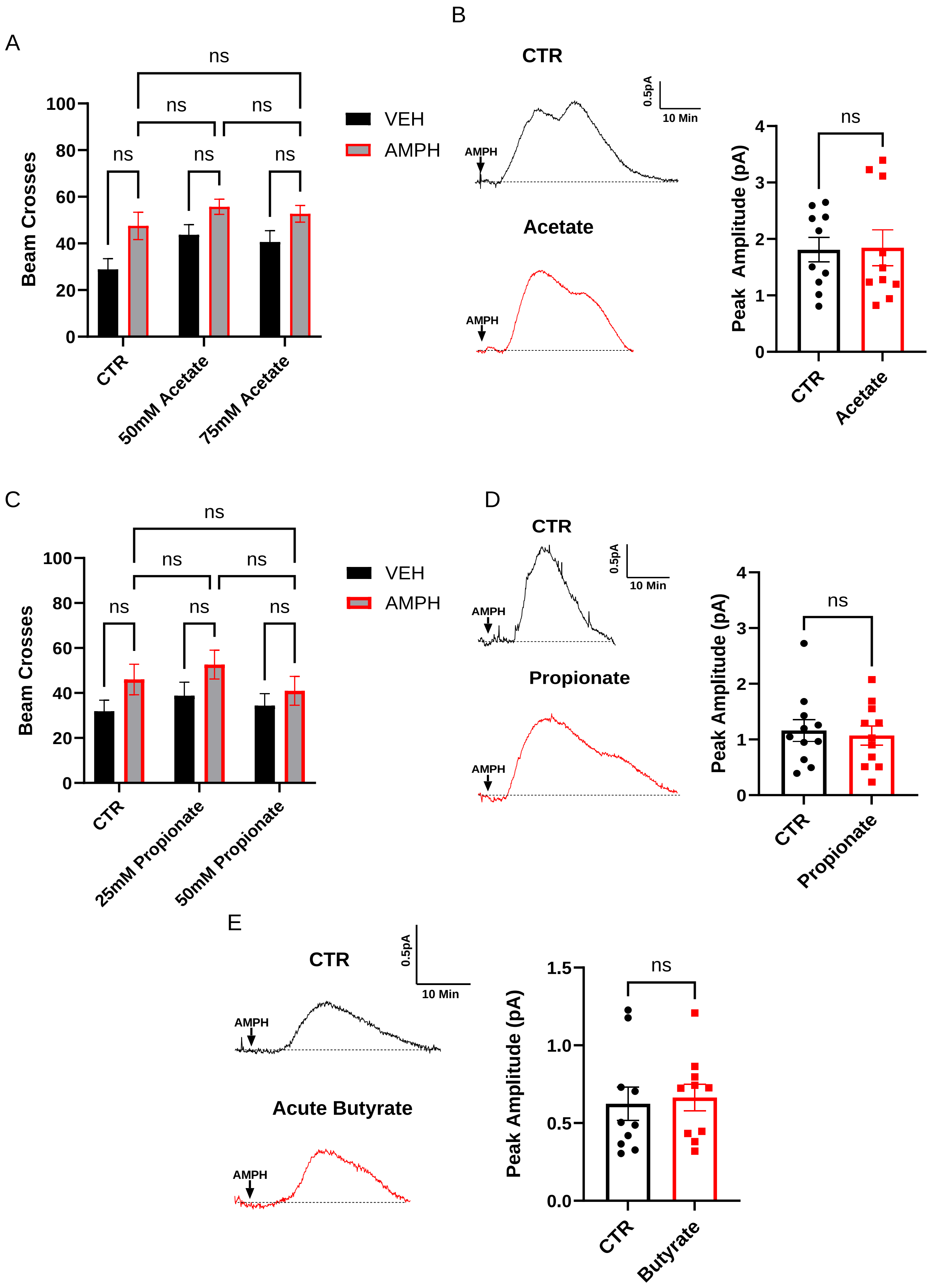

### Supplemental Figure 4

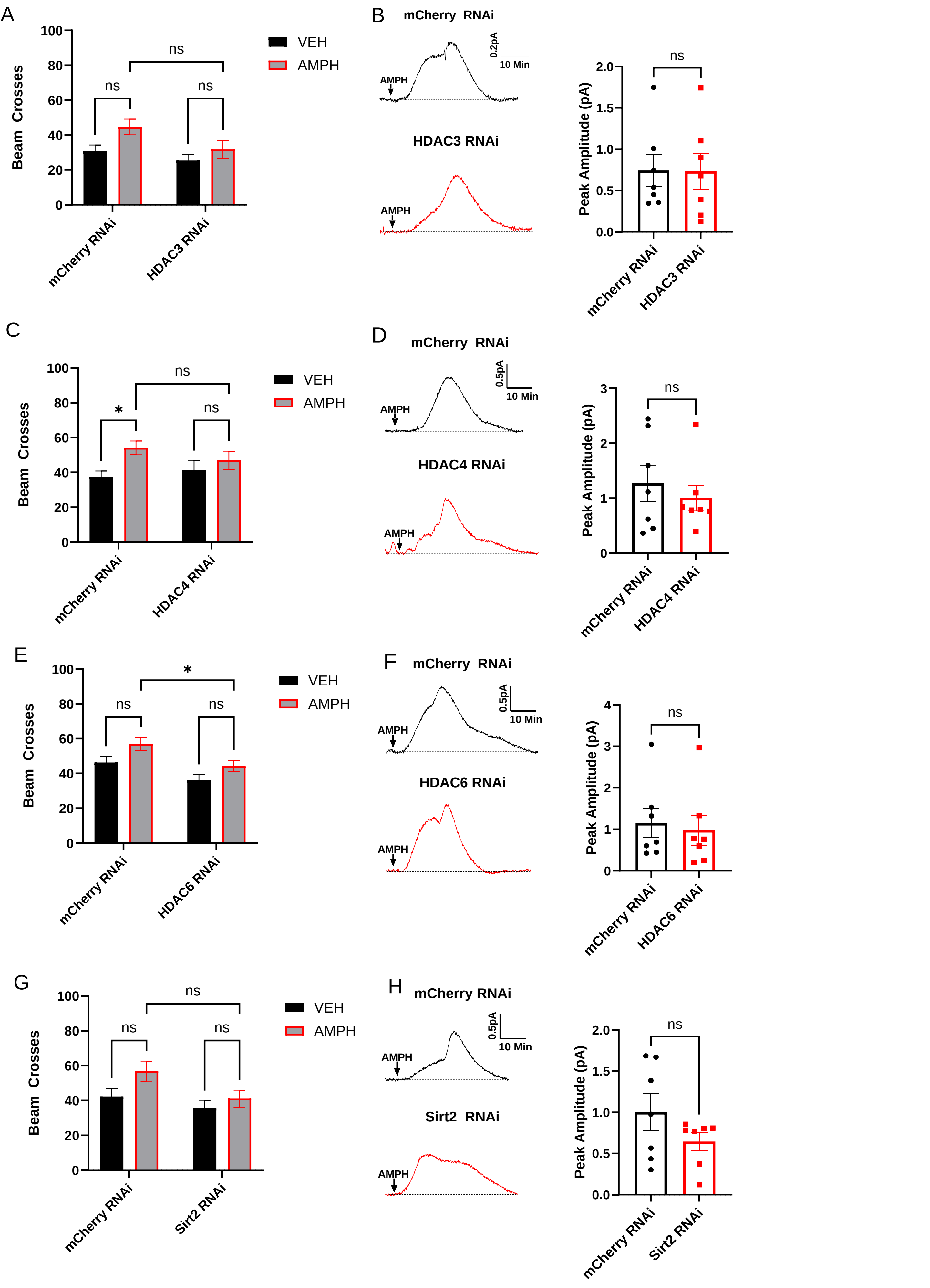

### Supplemental Figure 5

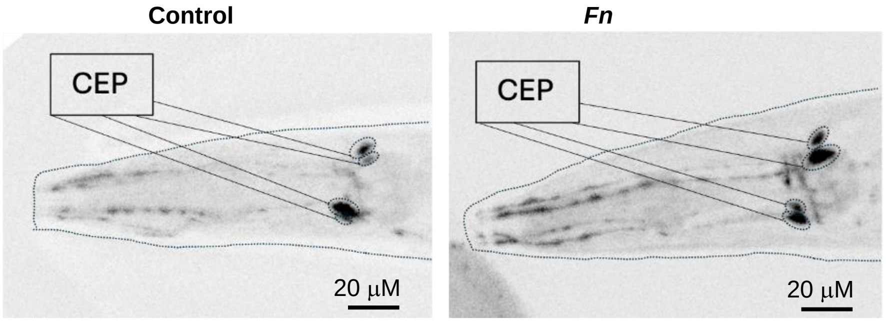

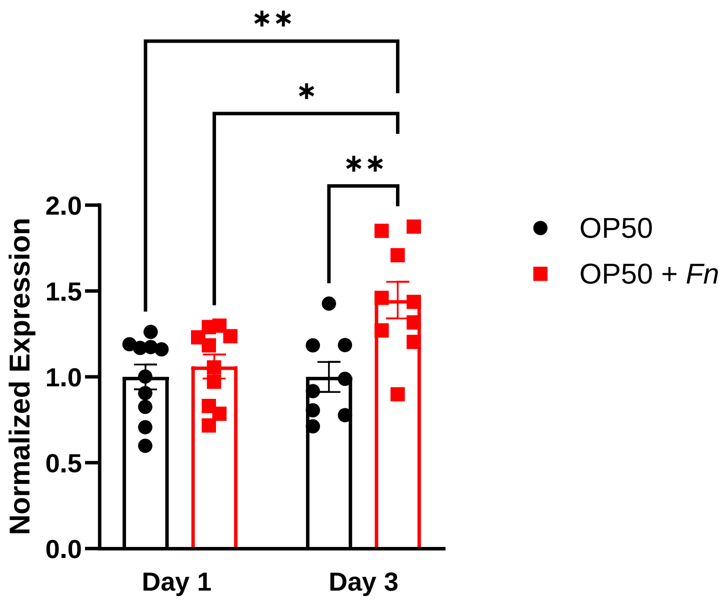

### Supplemental Figure 6

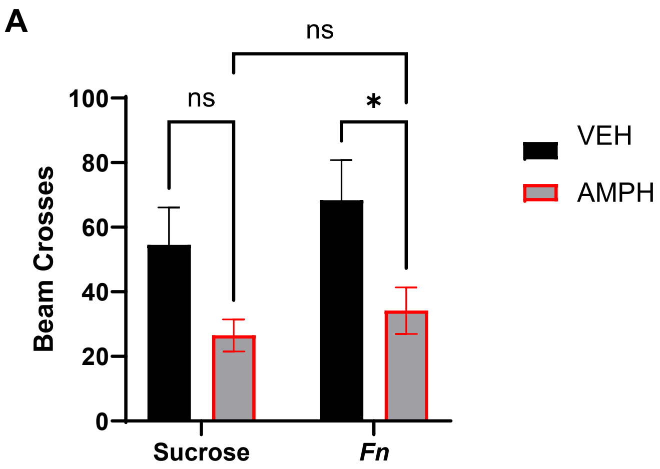

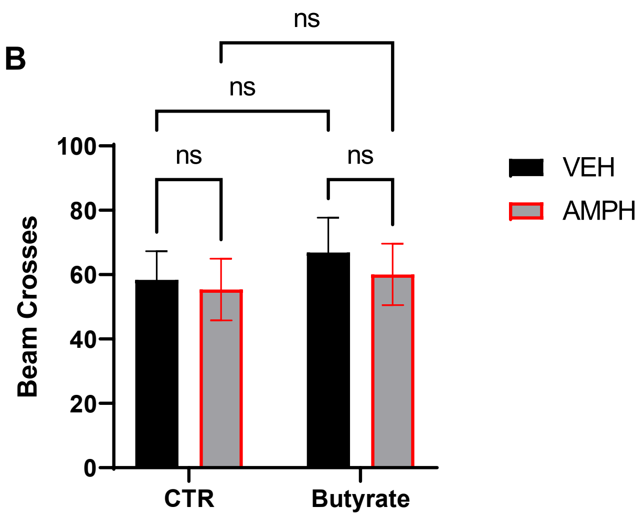

### Supplemental Figure 7

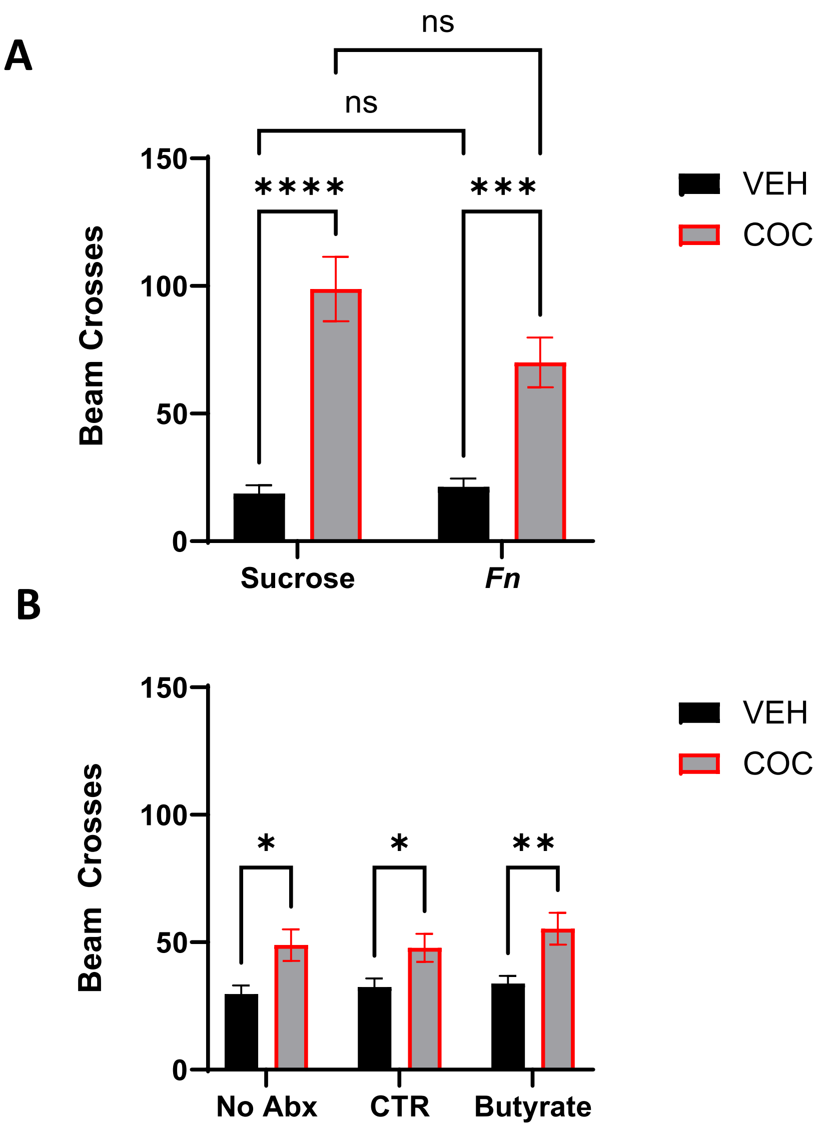

### Supplemental Figure 8

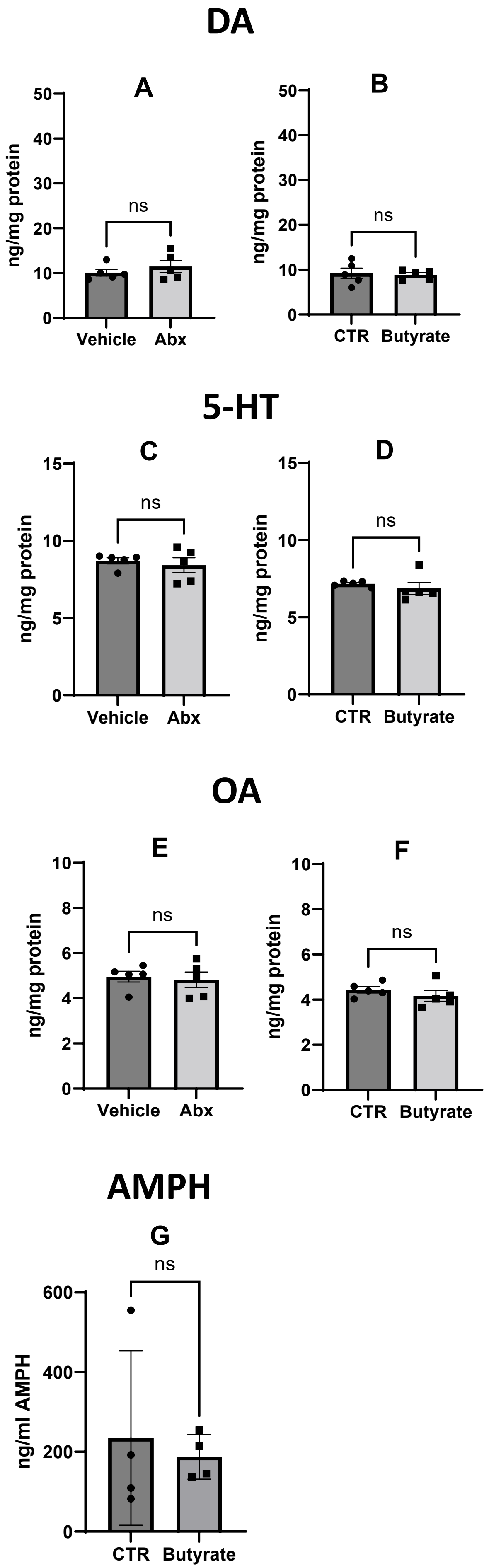

### Supplemental Figure 9

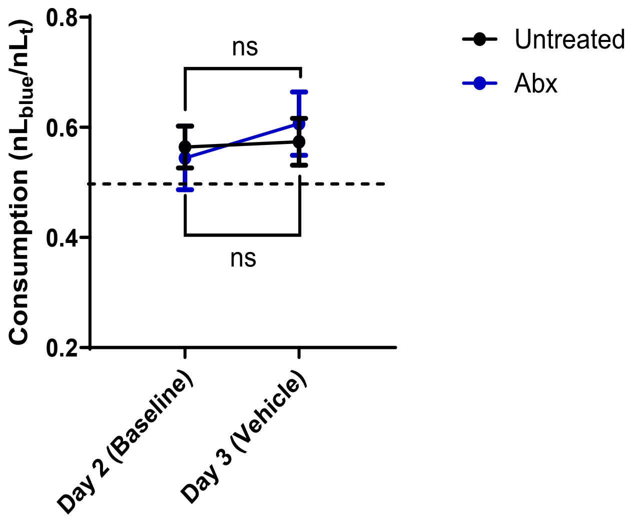

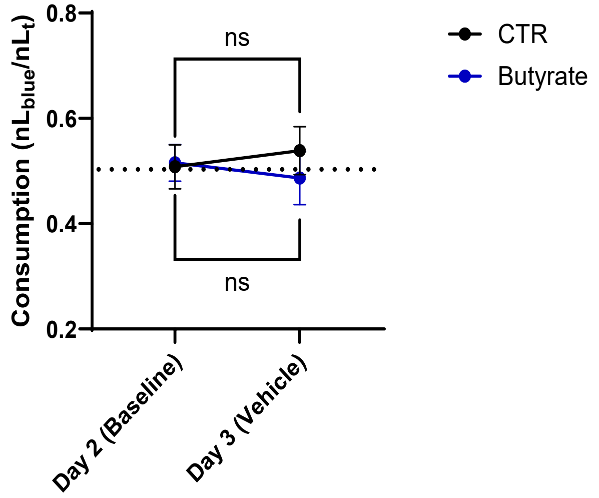

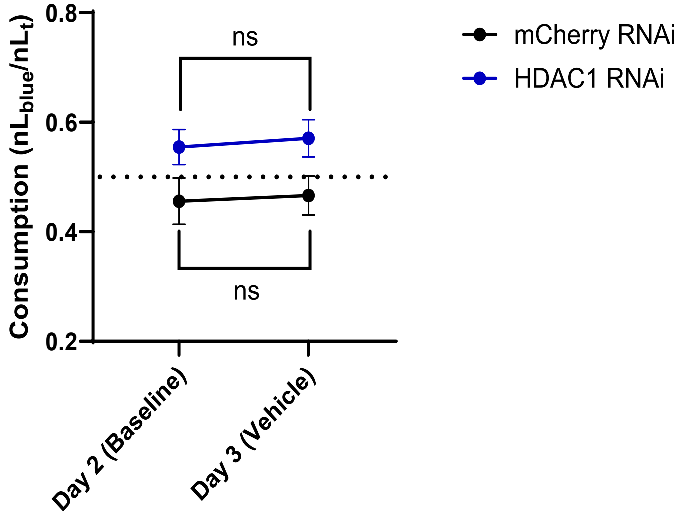
